# Neural circuits underlying context-dependent competition between defensive actions in *Drosophila* larva

**DOI:** 10.1101/2023.12.24.573276

**Authors:** Maxime Lehman, Chloé Barré, Md Amit Hasan, Benjamin Flament, Sandra Autran, Neena Dhiman, Peter Soba, Jean-Baptiste Masson, Tihana Jovanic

## Abstract

To ensure their survival, animals must be able to respond adaptively to threats within their environment. However, the precise neural circuit mechanisms that underlie such flexible defensive behaviors remain poorly understood. Using neuronal manipulations, machine-learning-based behavioral detection, Electron Microscopy (EM) connectomics and calcium imaging in *Drosophila* larva, we have mapped the second-order interneurons differentially involved in the competition between different defensive actions and the main pathways to the motor side putatively involved in inhibiting startle-type behaviors and promoting escape behaviors in a context dependent manner. We found that mechanosensory stimulation modulates the nociceptive escape sequences and inhibits C-shape bends and Rolls in favor of startle-like behaviors. This suggests a competition between mechanosensory-induced startle responses and escape behaviors. Structural and functional connectivity revealed that the second order interneurons receive their main input from projection neurons that integrate mechanosensory and nociceptive stimuli. The analysis of their postsynaptic connectivity in EM revealed that they make indirect connections to the pre-motor and motor neurons. Finally, we identify a pair of descending neurons that could promote modulate the escape sequence and promote startle behaviors. Altogether, these results characterize the pathways involved in the Startle and Escape competition, modulated by the sensory context.

## Introduction

It is essential for survival across the animal kingdom to avoid dangers and threats successfully. Different defensive behavioral strategies lead to appropriate avoidance of threats depending on their nature and properties^1–4^. The defensive strategies animals employ to evade potential dangers may include various modes of locomotion to flee from the perceived threat, as well as immobilization or freezing behaviors aimed at impeding detection by the aggressor. Additionally, animals may execute protective actions designed to minimize the exposure of sensitive areas^5^.The type of behavior that will be performed will depend on the type of danger as well as on the specific context in which the danger is encountered.

While some defensive behaviors are stereotyped to ensure rapid responses to threatening stimuli, they also need to be flexible as animals respond differently to threats depending on the environment and their internal needs^5–8^, with trade-offs and costs that take into account all these factors. Escape behaviors range from very simple reflex-like actions^9–11^ to more complex ones requiring cognitive processes relying on memory and decision-making^5,12,13^. Animals must first decide whether to respond and then how to respond to threats. Both decisions could be affected by the type and degree of threat but also context, internal or behavioral state. For example, hungry crayfish rather freeze than perform a tail flip in response to looming stimuli^14^, and feeding leeches ignore mechanical stimuli due to the blocking of the transmission of mechanosensory information to central circuits through presynaptic inhibition of mechanosensory terminals^15^. In mice the spatial environment will determine whether they will freeze or escape to shelter the location of which they memorize^16^. Moreover, defensive behaviors are often not single actions but multiple actions that need to be organized in sequences^5,6,17–19^, and the order of which may also vary depending on the context or state.

The neural circuit mechanisms underlying the flexibility of defensive behaviors, the selection between various competing actions, and their sequential organization based on context remain to be fully understood. We address this question in the *Drosophila melanogaster* larva, which due to its genetic tractability and the availability of the electron microscopy (EM) connectome^18^ of its 10 000-neuron central nervous system (CNS) allows the mapping of circuits at a synaptic and cellular level. The ease of circuit mapping, combined with the rapid life cycle and the ease of automated and quantitative behavioral approaches makes it an excellent model to relate neural circuit structure and function underlying defensive behaviors^20^.

To avoid and escape various threats, larvae perform different actions and use different strategies depending on the nature and degree of threat. For example, we have described previously using machine-learning-based algorithms that automatically detect larval actions^19^, that the air puff, an aversive stimulus, can trigger five different types of actions that consist of startle-like actions (Hunch and Stop) and Avoidance or Escape actions (Back-up, Crawl, and Bend) that can be organized in a sequence^17,19^. In response to nociceptive stimuli, larvae perform escape behaviors consisting of C-shape and Rolls^18,21–23^.

In this study, we identify second order interneurons that underlie the inhibition of startle response while at the same time promoting an escape sequence. We further find that the sensory context modulates the escape sequence induced by optogenetic activation of a specific set of neurons downstream of the previously described startle and escape circuit^17,19^. Modulation of the behavioral output depends on the stimulation strength of the 2^nd^ order interneurons and the mechanosensory context introduced by using an air puff. Altogether, our work reveals that the identified neurons can gate specific responses depending on the context and activation strength, contributing to the circuit computation for selecting appropriate defensive behavior.

## Results

### EM connectivity analysis reveals candidate sensorimotor neurons for avoidance responses to a mechanical stimulus

We have previously characterized the behavioral response of larvae to a mechanical stimulus, the air puff. We found that, when given air puff stimulation, larvae performed a probabilistic sequence of five different avoidance actions: Hunch, Bend, Back-up, Stop, and Crawl, and identified the pathways in the mechanosensory network that underlies these behaviors^19^. We have also described in detail the circuit for the selection between the two most prominent actions in the sequence: Hunching (head-retraction) and Bending^17^. The circuit is composed of chordotonal mechanosensory neurons that sense the air puff, a layer of different types of inhibitory neurons, and a layer of two projection neurons: Basin-1 and Basin-2 which receive inputs from chordotonal sensory neurons. The model of that circuit predicts that the coactivation of both projection neurons gives rise to a Bend, while the activation of Basin-1 would result in Hunching. Effectively, this suggests that activation of Basin-2 inhibits the Hunch. To determine how the information is decoded downstream of Basin neurons to give rise to Hunching or Bending, we looked in the EM connectome for candidate neurons downstream of Basins that could be involved in Hunching or Bending. One candidate neuron is A19c, previously identified in a behavioral screen labeled by the genetic driver R11A07^19^. The R11A07 also labels a thoracic neuron with descending projection (Fig. 5). In that screen, driving Tetanus-toxin (TNT) with the R11A07 driver resulted in an increase in Hunching probability in response to air puff, suggesting A19c inhibits the Hunch. Interestingly, A19c receives inputs from Basin-2 and Basin-4 neurons^18,19^, (Fig. 1C,D, Supplementary fig. 1A, B, Supplementary table 1). Basin-4 neurons as Basin-2, also inhibit the Hunch response^17^. Another group of candidate neurons downstream of Basins that could be involved in air puff responses are the interneurons A08m and A08x, which receive the largest fraction of inputs from Basin-1 and Basin-3 (Fig. C,E, Supplementary fig. 1C,D, Supplementary table 2). A08m receives the majority of its Basin inputs from both Basin-1 and Basin-3, with slightly more inputs from Basin-3 inputs than Basin-1 inputs in the first abdominal segment (Fig. 1E1, Supplementary fig. 1C, Supplementary table 2). A08x receives most of its Basin inputs from Basin-1 (in all segments where Basins were reconstructed) (Fig. 1E2, Supplementary fig. 1D). A08x and A08m neurons are part of a group of five neurons called early-born Even-skipped expressing lateral neurons (ELs)^24,25^. These neurons respond to mechanical stimulation and their optogenetic activation was shown to induce escape Rolling^25^.

**Figure 1.**
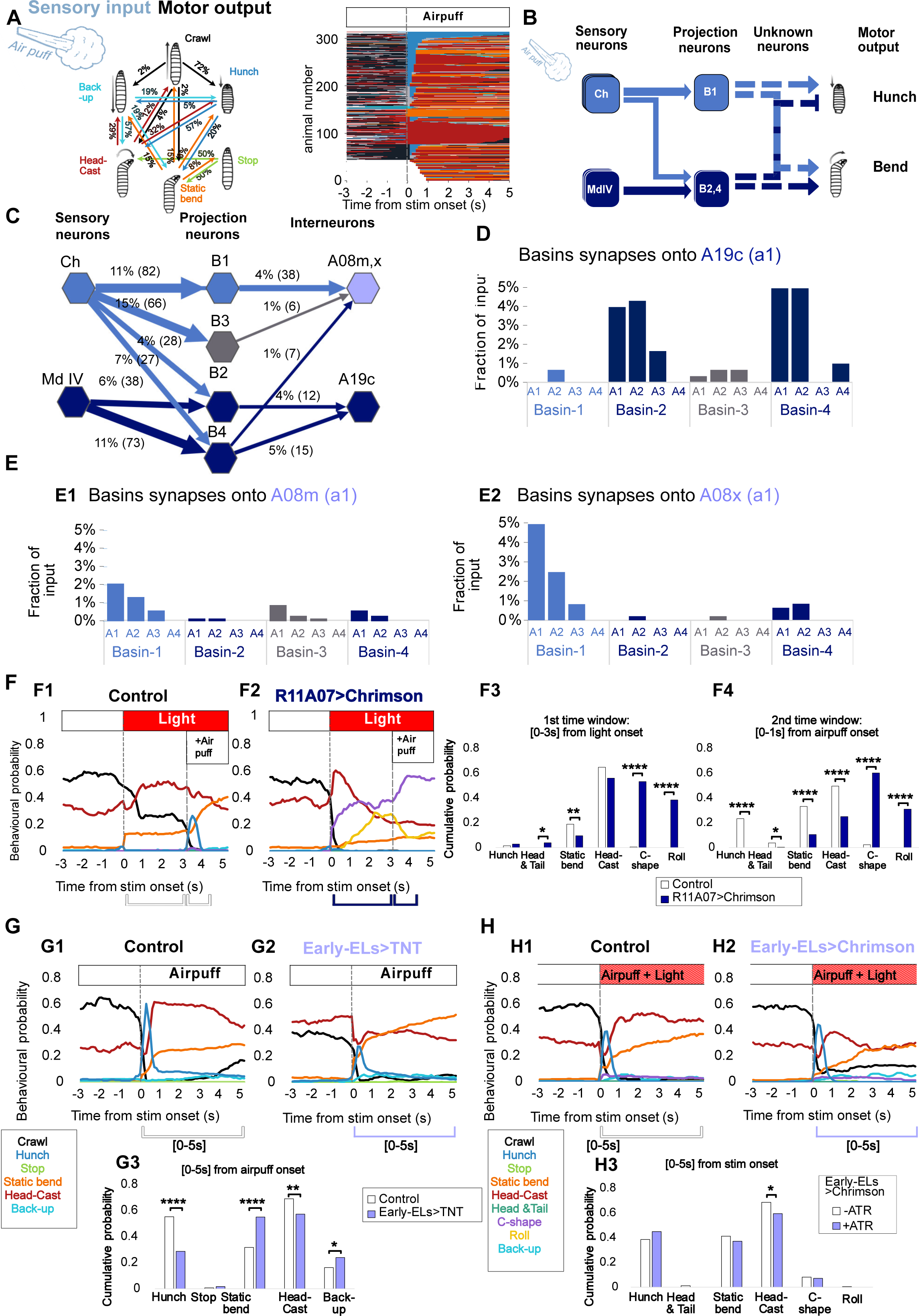
Second order interneurons in a mechanosensory network are differentially involved in responses to air-puff. **A.**Drosophila larvae respond to a mechanosensory input, the air puff, with probabilistic sequences of actions <u>(Masson et al., 2020)</u>. Left: schematic showing behavioral probabilities of immediate response to an air puff and of action transition (from <u>(Masson et al., 2020)</u>. Right: ethogram showing larval behavioral responses of across time, with one line representing one individual and one color representing one action as in A. Control genotype: attP2>TNT (n=818). The ethogram was organized to ease reading by grouping larvae by their first behavioral response to stimuli. **B.** Simplified schematic of the previously characterized circuitry underlying Hunch and Bend responses to air puff <u>(Jovanic</u> <u>et al., 2016)</u>. Ch represents chordotonal mechanosensory neurons, Md IV represents nociceptive class IV multidendritic neurons, B1 represents Basin-1, B2,4 represents Basin-2 and Basin-4. **C.** Graph showing synaptic connectivity based on EM reconstruction between neurons depicted in B, as well as the second-order interneurons A08m,x and A19c. Connections were shown as follows: fraction of total input (number of synapses). In light blue are all neurons previously identified as being required for Hunch, in dark blue are all neurons previously identified as inhibiting Hunch. B3 is shown in gray as its role in hunching is unknown. Only neurons in neuromere a1 are shown. **D.** Input A19c receives from Basins across neuromeres where Basins were reconstructed: a1-4, shown as percentage of all input A19c receives **E.** Input A08m and A08x receive from Basins across neuromeres a1-4,Connections represented as a of all input A08m receives (in a1 where it was reconstructed) and fraction of all input A08x receives (in a1)). **F.** Larval responses to light (at 60s.) then air puff, delivered 3s after the light activation. **F1** control larvae (attP2>CsChrimson, n=177) **F2.** larvae with activated R11A07 neurons (R11A07>-CsChrimson, n=250). **F1-F2.** mean behavioral probability over time **F3.** behavioral probability cumulated over the first three seconds after light onset, corresponding to control larvae (white) and larvae with R11A07 activated (dark blue) **F4.** behavioral probability cumulated over the first second after air puff onset, corresponding to control larvae (white) and larvae with R11A07 activated (dark blue). **G.** Larval responses to 4m/s air puff. **G1.** control larvae (eELGAL4>CantonS n=254) **G2.** larvae with Early-ELs interneurons inactivated (eEL- Gal4>TNT, n=420). **G1-2.** mean behavioral probability over time. Stim onset at 60 s. **G3.** behavioral probability cumulated over the first five seconds after air puff onset, corresponding to control larvae (white) and larvae with early-ELs inactivated (lavender). **H.** Larval responses to air puff (4m/s intensity) and optogenetic activation (0.3mW/cm² irradiance). **H1**, control larvae (eEL-Gal4>CsChrimson, without ATR n=248), **H2.** larvae with Early-ELs interneurons expressing CsChrimson (eEL-Gal4>CsChrimson, with ATR n=286). **H1-H2** mean behavioral probability over time. Stim onset at 60 s **G3.** behavioral probability cumulated over the first five seconds after air puff onset, corresponding to control larvae (white) and larvae with early-ELs activated (lavender). For all barplots: *:p<0.05, **:p<0.005, ***:p<0.0005,****:p<0.0001, Chi² test

### Second-order interneurons in a mechanosensory network are differentially involved in Startle and Escape responses to air puff

To determine whether A19c and A08m,x could be involved in decoding the Basin-1 and Basin-2 activities to induce appropriate motor outputs, we investigated the role of A19c and early-born ELs in air-puff-induced sensorimotor decisions. For that purpose, we employed high-throughput behavioral assays that we have previously established^17,19,26^ and an updated classification method that we have developed in this study. The new classifiers disentangle two types of Bend behaviors: Head Casting and Static Bending, in addition to detecting Hunch, Stop, Back-up, Crawl, and Roll (Fig. 1A). Head Casting consists of the larva swiping its head to explore the environment once or multiple times. It occurs both in the presence and absence of sensory stimulation in the context of, for example, navigating sensory gradients or foraging respectively^27–35^.Using the new classification method, the finding from the screen where driving TNT with the R11A07 driver resulted in an increased Hunching probability in response to air puff was confirmed (Supplementary fig. 2A)^19^.To observe whether the optogenetic activation would be sufficient to inhibit Hunching, the same driver was used to express CsChrimson in these neurons and optogenetic responses were induced with red light while simultaneously delivering air puff (Supplementary fig. 1B). Indeed, optogenetic activation of neurons labeled by the R11A07 driver resulted in a decrease in Hunching as well as Static Bending and Head Casting (Supplementary fig. 2B). Moreover, if air puff was delivered 3 s after the light onset, no Hunching at all was observed (Fig. 1F). Interestingly, R11A07>CsChrimson optogenetic activation triggered an escape response involving C-shapes and Rolling (Fig. 1F and Supplementary fig. 2B), typically elicited in response to noxious stimuli.

We then analyzed all the inputs to the A19c using EM connectomics^36^ and found that Basin-2 and Basin-4 represent nearly a quarter of all of the A19c inputs and no significant input from Basin-1 was observed (Fig. 1C,D, Supplementary fig. 1, Supplementary table 1). We previously showed that Basin-4, similarly to Basin-2, inhibits Hunching and is required for Bending^17^. In addition, it has been shown that the optogenetic activation of Basins triggers Rolling^18^. Since silencing A19c resulted in a similar behavioral phenotype to the phenotype of its main presynaptic partners, Basins-2 and ™4, this could suggest that Basins-2 and ™4 could inhibit Hunch and promote escape behaviors (Rolling) through A19c.

We further investigated the behavioral role of early-born ELs, by silencing them during air puff responses using TNT and found that they are required for Hunching (and Head Casting) (Fig. 1G, Supplementary fig. 3C,D) and inhibited Static Bending and Backing-up as silencing resulted in less Hunching and Head casting and more Static Bending and Back-ups. Furthermore, optogenetic activation of early-ELs during air puff resulted in increased Hunch probability in response to air puff (Fig. 1H, Supplementary fig. 3A,B), showing that early-ELs promote Hunching, as do their presynaptic partners Basin-1. However, previous work showed that silencing of Basin-1 results in both less Hunching and less Bending. Conversely, silencing of Els results in more Static Bending. It is important to note that the inactivation and the optogenetic activation experiments involved all five early-born ELs (as targeted by the genetic line EL- Gal4,R11F02-Gal80) and that out of these five neurons, only A08m and A08x receive significant input from Basins. Analyzing A08m,x presynaptic connectivity in the EM volume, we found that these neurons received a significant percentage of their input from Basin-1 (and ™3) and a relatively smaller percentage of input from Basins 2 and 4 (Fig. 1E, Supplementary fig. 1C,D). Interestingly, optogenetic activation of early-ELs was previously described to trigger a Rolling escape response^25^, which we also found at high optogenetic intensity (2.2mW/cm² irradiance), and could be explained either by the input A08m,x received from Basin-2 and ™4, or by the implication of the other three neurons in escape behaviors Together, these results suggest that A19c and early-ELs A08m,x are second-order interneurons that could be differently involved in Hunching and Escape behaviors. The early ELs, downstream of Basin-1 seem to be required for Hunching and inhibit Static Bending, while the A19c, downstream of Basin-2 and ™4 could inhibit Hunching (and Static Bending) and promote an escape sequence.

### Automated classification of Bend-like behaviors reveals stereotyped escape sequences

We further characterized in detail the escape sequence triggered by optogenetic activation with the R11A07 driver. By inspecting larval behavior recordings upon R11A07>CsChrimson optogenetic activation, we found that, as previously reported, in the Escape sequence, the Roll was preceded by a C-shape^23^. We also observed that some larvae initiated the Escape sequence with a symmetrical contraction that resembled a Hunch but consisted of both the Head-and-Tail contractions, before transitioning into a C-shape (Fig. 2). Indeed, the amplitude of larval length decreases is higher upon light stimulation which induces optogenetic activation with R11A07 than upon air puff that induces Hunch (with a smaller decrease in length compared to Head-and-Tail contraction) (Supplementary fig. 4A). This behavior was not thus far described and has not been observed as being part of the Rolling Escape sequence.

**Figure 2.**
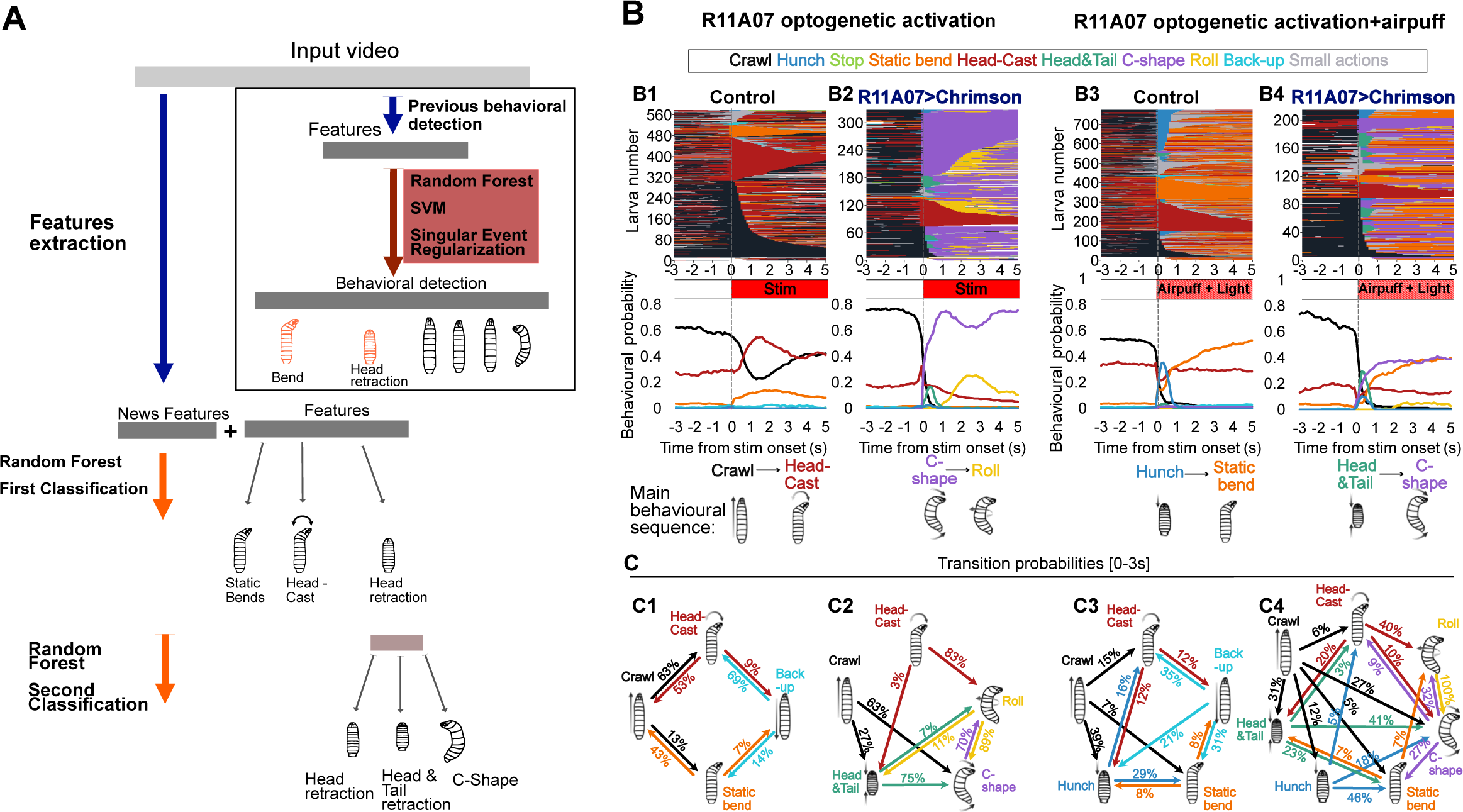
An Automated classification of Bend-like behaviors reveals stereotyped escape sequences triggered by optogenetic activation of R11A07 neurons. **A.**Reclassification procedure of larval behaviors automatically detected by a Machine Learning-based algorithm. Actions previously categorized as “Bend’’ were reclassified to be categorized as either “Head-Cast”,”Static Bend’’ or “C-shape”, and actions previously categorized as “Hunch”(head-retraction) were reclassified to be categorized as either “Hunch”,“Head&Tail” or “C-shape”. **B.** Larval responses to **B1** only light (control: attP2>CsChrimson, n=530) **B2.** only light (optogenetic activation, R11A07>CsChrimson, n=305). **B3.** air puff and light (control: attP2>CsChrimson ATR,n=765) **B4** air puff and light (optogenetic activation, R11A07> CsCrimsonn=214) Top: ethograms, actions over time, with one line corresponding to one individual and each color corresponding to a different action (as indicated). Bottom: mean behavioral probability over time. Stim onset at 0s. air puff intensity: 3 m/s. Irradiance: 0.3mW/cm² C. Schematic depicts the characteristic behavioral sequence for each condition **D.** transition probabilities cumulated over the first three seconds after stim onset. Only transition probabilities of 3% or more are shown **D1**: Control, light alone. **D2**: R11A07c>CsCrimson, light alone. **D3**: Control, air puff and light. **D4**: R11A07>Cs Crimson, air puff and light.

To quantify the probability of the different actions in the Escape sequence in different conditions of stimulation, we worked to further expand the previously developed machine learning-based behavioral classification algorithms^19^ to also detect Head-and-Tail and C-shape actions. We thus adapted the classification such that what was previously categorized as “Bend” would now be either classified as “Head Cast”, “Static Bend” or “C-shape” and what was previously categorized as “Hunch” (Head-retraction) would now be classified as either “Hunch”, “Head-and-Tail” or “C-shape” (Fig. 2A). The new classification is computed with a simple Random Forest algorithm based on a new set of features and newly annotated data (see detailed description in the Method section).

Using the new classification algorithms, we examined the behavior sequence in response to an air puff, optogenetic activation of R11A07 neurons, or both (Fig. 2). We found that larvae predominantly responded to an air puff (in combination with red light) with either a Hunch, a Static Bend, or a Head Cast (Fig. 2B3). Hunch and Static Bend are indeed not seen before the stimulus, while Head casts can be observed even in the absence of stimuli as larvae typically explore their environment by Crawling interspersed with Head casts and Turns^35,37^. The different behavioral actions that larvae perform in response to various somatosensory stimuli can be categorized into, on the one hand, static, startle-like actions like Stopping, Hunching, and Static Bending and, on the other, active-type, exploratory and escape actions (Crawling, Head cast, Fast Crawl, Head-and-Tail contractions, C-shape, and Roll)^17–19,22,23,26,38^. Upon optogenetic activation of R11A07 alone, most larvae perform a C-shape (63% out of larvae crawling) that is sometimes followed by a Roll (Fig. 2B2,C2, Supplementary table 3). A smaller percentage of larvae will perform a Head-and-Tail contraction (27% of Larvae that were crawling) followed by a C-shape (Fig. 2B2,C2, Supplementary table 3). C-shapes are in 70% of cases followed by a Roll. If optogenetic activation was done in combination with air puff, the larvae also performed Hunches and Static Bends. Out of the escape actions (Head-and-Tail, C-shape and Roll), they perform relatively more Head-and-Tails (30%) than when optogenetic activation was done alone and fewer larvae (45% compared to 67% upon optogenetic activation alone) perform C-shapes (out of the larvae that were crawling prior to the stimulus) (Fig. 2B4,C4, Supplementary table 3). In most cases (69%) Head-and-Tail contractions were followed by a C-shape. C-shapes are less frequently followed by a Roll (30%) than when the light was delivered alone (Fig. 2B-D, Supplementary table 3). These results reveal that different types of escape sequences are induced by R11A07>CsChrimson activation depending on whether it was coupled with mechanical stimulation or not: a Head-and-Tail > C-shape sequence is more prominent when the activation is coupled with air puff, while the C-shape > Rolling sequence predominantly occurs upon optogenetic activation alone (Fig. 2B-D). These results raise the question of how the sensory context shapes the dynamics of nociceptive escape sequences.

### Sensory context modulates the optogenetically triggered escape sequence

To investigate the influence of the mechanosensory stimulation on Escape sequences triggered by optogenetic activation using the R11A07 driver, we subjected R11A07>CsChrimson larvae to different intensities of air puff and light stimulation (Fig. 3, Supplementary fig. 5).

**Figure 3.**
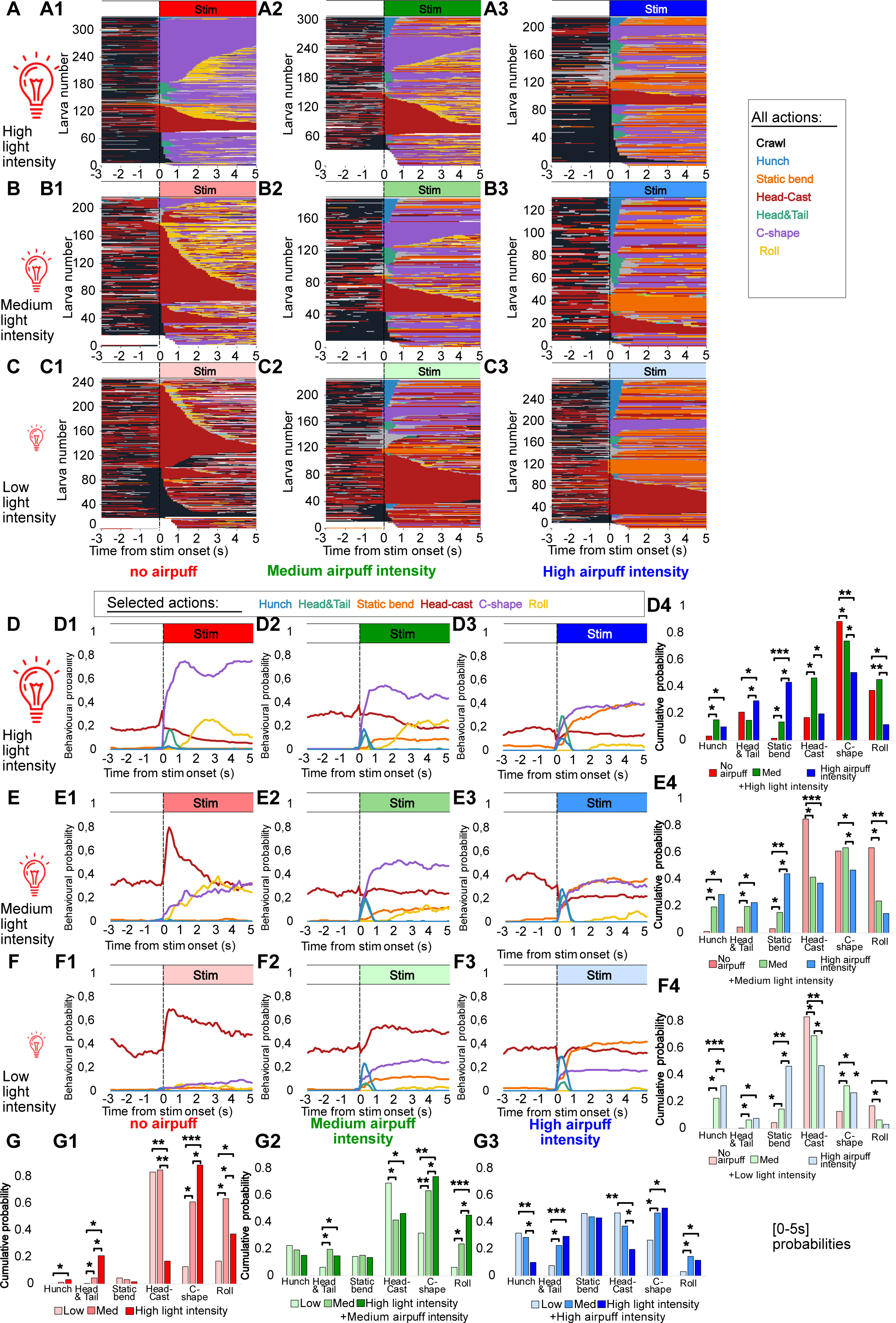
Sensory context modulates escape responses triggered by R11A07>CsCrimson optogenetic activation. **A-C.**Ethograms, corresponding to the same experiments and organized in an identical matrix as that shown above (A-C) including all detected actions. **D-F.** behavioral probabilities of Hunch, Head&Tail C-shape, Roll, in response to: **D.** high light intensity (0.3mW/cm² irradiance) and various intensities of air puff: **D1**: no air puff (n=305). **D2**: 3m/s air puff (n=265). **D3**: 4m/s air puff (n=214) **E.** medium light intensity (0.2mW/cm²) with **E1**. no air puff (n=183) **E2**. 3m/s air puff (n=175). **E3**. 4m/s air puff (n=129). **F.** low light intensity (0.1mW/cm²) with **F1**: no air puff (n=199). **F2**: 3m/s air puff (n=206). **F3**: 4m/s air puff (n=269). **D4, E4, F4, G1-3.** Barplots correspond to the behavioral probability cumulated over the first five seconds after stim onset. For all barplots: FDR: *:p<0.05, **:p<0.01, ***:p<0.005 Chi² test, with Benjamini-Hochberg correction

In the absence of air puff, we observe distinct Escape sequences depending on the level of irradiance used for optogenetic activation (Fig. 3): Head Cast and Crawling are prominent at low light intensity (0.1mW/cm²), with very little C-shape or Rolling (following Head Casts) (Fig. 3C,F,G). Although Head Cast and Crawl occur in response to light stimulation, the R11A07>CsChrimson larvae showed increased Head Casting and decreased Crawling compared to the control upon light stimulation (Supplementary fig. 5). The increase in Head Casting could be interpreted as mild escape behavior triggered by a low activation level. Medium-high light intensity (0.2mW/cm²) resulted in a stronger escape response, with significantly more C-shapes and Rolls, at the expense of Crawling (Fig. 3B,E,G, Supplementary fig. 5B2). In addition, more larvae transitioned from a Head Cast into a Roll (Supplementary fig. 5B3,C3). Increasing light intensity even more (increased the transition probabilities from Head Cast into Roll even more and drastically increased the C-shape probability at the expense of Head Casting and, surprisingly, of Rolling (Fig. 3G1, Supplementary fig. 5A3,B3,C3). In addition, at high light intensity, the Head-and-Tail probability increased significantly (Fig. 3A,D,G). These results suggest that different levels of optogenetic activation result in various Escape sequences. Lower levels of optogenetic activation trigger mild escape responses composed of mainly Head Casts (sometimes followed by Crawls) and low levels of C-shapes and Rolls, while the medium and high levels of the activation trigger an escape sequence consisting of either C-shapes and Rolls or Head-and-Tail and C-shapes (and Rolls) respectively. The Crawls observed upon R11A07 optogenetic activation are faster compared to the control and therefore represent the Fast Crawl escape (Supplementary fig. 4B). Interestingly the strongest optogenetic activation of R11A07 neurons triggers a majority of C-shapes and not Rolls.

To determine how other types of sensory information will affect these escape sequences, we delivered air puff stimulation of two different intensities: medium and strong, along with different levels of optogenetic activation. The air puff appears to modulate the escape sequences triggered by the optogenetic activation: Rolling is inhibited by both medium and strong air puff at low and medium light intensities (optogenetic activation) (Fig. 3E4,F4). This decrease in Rolling probability is accompanied by an increase in Head-and-Tail contractions at both air puff intensities (Fig. 3E4,F4). At high levels of optogenetic activation, only upon delivering strong air puff can we see an increase in Head-and-Tail probability at the expense of Rolling (Fig. 3D4). This intensity-dependent increase in Head-and-Tail contractions and decrease in Rolling in the presence of air puff stimulation suggest context-dependent competitive interactions between different actions within the escape sequence.

Similarly, at the medium air puff intensity, an increase in C-shape probability at the expense of Rolling can be observed at low and medium light intensity (Fig. 3G2). This increase in C-shapes is weaker when the air puff intensity is higher at low levels of optogenetic activation (Fig. 3G3). At high air puff intensities and medium levels of optogenetic activation, C-shapes are decreased in favor of Head-and-Tails but also Hunches and Static Bends (Fig. 3D4,G3). Thus, in addition to the competition between different actions of the escape sequence, the presence of air puff that triggers Static Bend and Hunch during optogenetic activation induces the competition between escape-type and startle-type actions. Static Bend and Hunch probabilities increase with air puff intensity at low and medium optogenetic activation (Fig. 3B,C,E,F,G). At high levels of optogenetic activation, the Static Bend probability is increased at both air puff intensities, while Hunch is increased only at medium air puff intensity (at the expense of Head-and-Tail contractions) (the increase in Static Bend being stronger at this air puff intensity) (Fig. 3A,D, Supplementary fig. 6B). Altogether these data suggest that there could be competition between mechanosensory-induced Startle response and Escape behaviors triggered by optogenetic activation of the R11A07 neurons.

The finding that the different competitions (between the different actions in the sequence and between escape and startle behaviors) occur depending on the relative level of activation of R11A07 neurons and the air puff stimulus intensity, suggests that the competitions occur at the level of these neurons or downstream of them.

### R11A07 neurons are required for mechano-nociception

To determine whether the A19c could be also required for nociceptive behaviors we inactivated the R11A07 neurons with an inwardly rectifying potassium channel Kir2.1 while delivering nociceptive stimuli and monitored larval behaviors. Upon application of a mechano-nociceptive stimulus with a 50 mN calibrated *von Frey* filament, larvae perform either a C-shape bend, a Roll or a Turn (non-response) (Fig. 4A). Larvae in which the R11A07 were silenced showed decreased nociceptive responses, and specifically Rolling compared to the genetic controls (Fig. 4A). In addition to nociceptive mechanical stimulation, Rolling has been shown to be triggered by local exposure to temperature above 40°C and this thermo-nociceptive Rolling is also mediated by multidendritic class IV neurons^21,39^. To determine whether the R11A07 neurons’ role in nociceptive behaviors extends to other modalities, we investigated larval thermo-nociceptive responses. We applied a thermo-nociceptive stimulus using a temperature controlled hot probe and measured larval Rolling response latencies. Silencing R11A07 neurons did not affect thermo-nociceptive responses. These results show that R11A07 neurons are required for mechano-nociceptive but not thermal-nociceptive behaviors.

**Figure 4.**
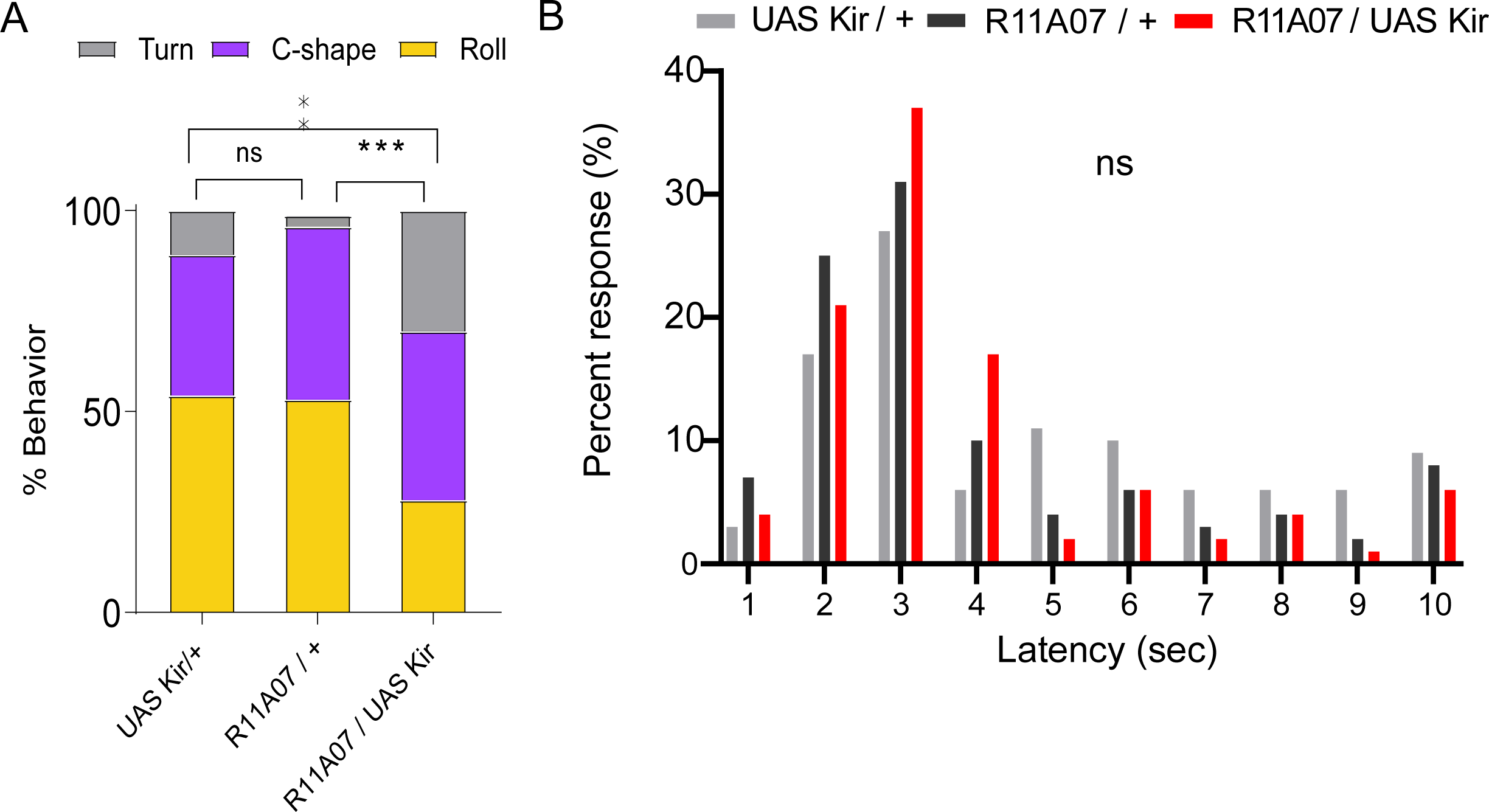
A-B. The R11A07 neurons are required for mechano-nociception but not thermo-nociception. **A.**Silencing R11A07 neurons using KIR results in less C-shape and rolling behavior in response to a mechano-nocipetive stimulus compared to the controls (p=0.0002, p=0.0049), n= 63 (UAS KIR/+), 60 (R11A07/+), 60 (R11A07/KIR) animals **B.** Silencing R11A07 didn’t affect latency of responses to a thermo-nociceptive stimulus, n= 70 (UAS KIR/+), 99 (R11A07/+), 90 (R11A07/KIR) animals Chi-square test was used to compare behavioral probabilities in A. **:< 0.01, ***: <0.001, Kruskal-Wallis test was used to compared latencies in B,

### Thoracic and abdominal neurons in the R11A07 line have distinct presynaptic connectivity and neurotransmitter identity

The R11A07 driver labels two types of neurons: abdominal A19c and a thoracic descending neuron (TDN) (Figure 5A). To better understand which of the neurons could be responsible for the phenotypes described thus far: triggering Escape sequences, inhibiting Hunching and Static Bending and taking part in context-dependent competitive interactions, between static actions (Hunch and Static Bend) and the escape sequence, we sought to investigate the presynaptic connectivity of these two neurons

**Figure 5.**
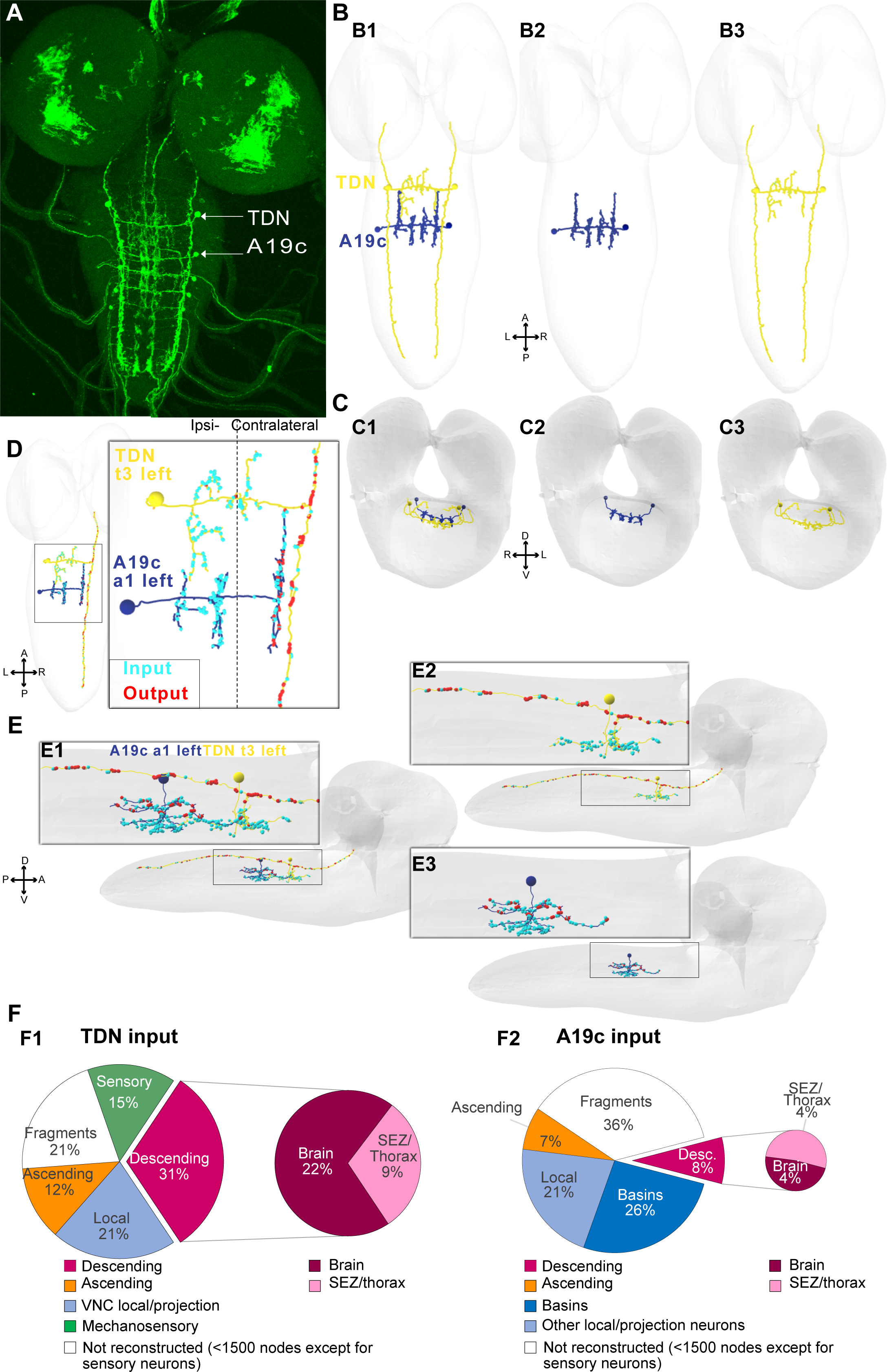
Morphological characterisation of TDN and A19c neurons. R11A07 expression pattern (GFP). thoracic neuron in the segment T3, and four abdominal neurons in segments a1-4. Elsewhere in the CNS, expressing GFP in this line, are bundles of cell bodies likely corresponding to immature neurons. **B.** Reconstructed A19c neuron in abdominal segment 1 (a1) (A19c, a1,left, right), in blue, and TDN, (t3, left,right) in yellow, antero-posterior view.**C:** Reconstructed A19c and putative TDN, dorso-ventral view. **D**. distribution of inputs(cyan) and outputs (red) in TDN_t3l and A19c_a1l, in an antero-posterior view. **E**. same as in D, viewed through a dorso-ventral angle. **F**. distribution of all input received by **F1**.TDN (t3l,t3r) **F2**. A19c (a1l,a1r). In white are all inputs from neurons not reconstructed up to recognition, i.e. fragments. We considered all neuron skeletons with fewer than 1500 neurons to be unreconstructed, except for sensory neurons (in green)

The morphology of the abdominal and thoracic neurons in the R11A07 line is very different: the thoracic neuron sends descending lateral projections spanning nearly the entire length of the ventral nerve cord (VNC). We identified in the EM images a neuron with morphological features similar to the TDN in the R11A07 driver line (Fig. 5A,B). The axon of the thoracic neuron is more lateral and more dorsal compared to the one of abdominal A19c (Fig. 5A-D). Both neurons have axonal projections that carry most of the output sites on the contralateral side with respect to the cell body. In contrast, the dendritic projections with most of the inputs are located on the ipsilateral side (Fig. 5C,D). The contralateral projections could play a role in controlling motor action requiring asymmetric contractions, such as Bends, C-shapes, and Rolls^24,40^.

We further analyzed all the input of the TDN neuron (Fig. 5F, Fig. 6.A,B, Fig. 4, Supplementary table 1). The analysis revealed that the TDN receives inputs from chordotonal subtypes lch5-2/4 in multiple neuromeres (a1, and a3-a8) (Fig. 6A,B, Supplementary table 1). This suggests that TDN could integrate mechanosensory input across segments. Analyzing the distribution of all presynaptic partners of the neuron, we found that this mechanosensory input represented 15% of all inputs. A significant fraction of inputs is from descending neurons (31 %), from local neurons in the VNC (21 %), and 12 % of inputs come from ascending neurons (Fig. 5F, Fig. 6A,B). Comparing the input distribution of TDN and A19c neurons revealed the difference in presynaptic connectivity between the two neurons (Fig. 5F, Fig. 6A,B). A19c receives most of its inputs from local and projections neurons in the VNC (47%), out of which Basins represent more than half of neurons. Only 8% are descending inputs, and A19c does not receive direct input from sensory neurons or ascending input (Fig. 5F, Fig. 6A,B, Supplementary table 1). On the contrary, TDN gets 15% of its inputs from sensory neurons and 12 % from ascending neurons. It also has more descending inputs (31%) than A19c. It is worth noting that most of the descending input TDN receives from Brain and SEZ neurons consist of axo-axonic connections (Fig. 5B, Supplementary table 1) along the VNC, suggesting that higher-order neurons could modulate the output of the TDN neurons depending on context, state, or experience.

**Figure 6.**
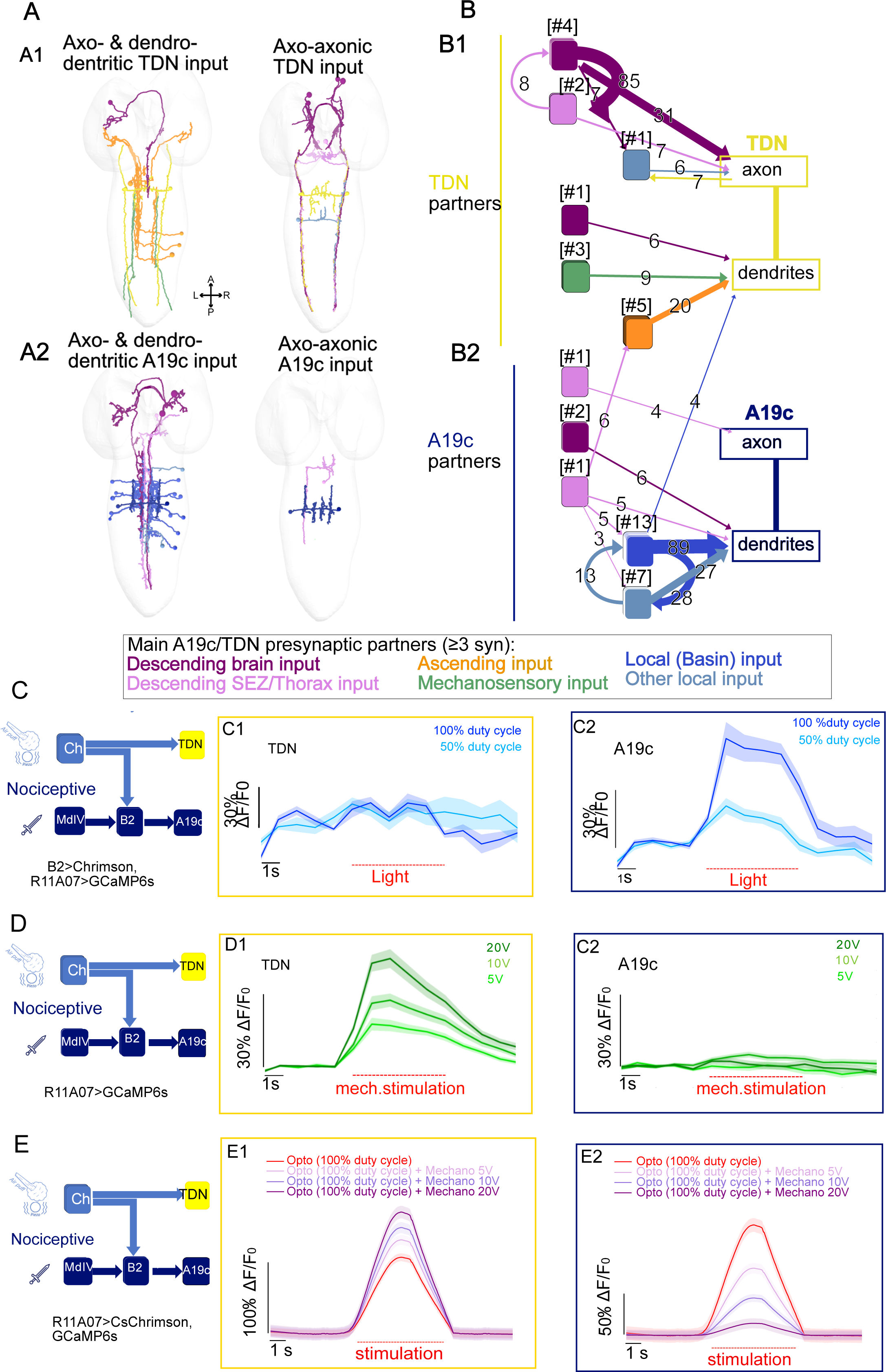
TDN and A19c neurons receive distinct inputs. **A, B.**Main presynaptic partners (with 3 or more synapses) of TDN, A19c. Fragments were not included. **A.** reconstructed skeletons shown in EM volume, using web-based software Catmaid. **A1**. TDN and its presynaptic partners, which send axo- and dendro-dendritic synapses (left), and axo-axonic synapses (right) onto TDN **A2**. A19c and its presynaptic partners, which send axo- and dendro-dendritic synapses (left), and axo-axonic synapses (right) onto A19c. Presynaptic partners were coloured based on their localisation (brain, subesophageal zone (SEZ), sensory (cho), local, ascending neurons. **B.** Connectivity between neurons shown in **A.** All connections of 3 or more synapses are shown. **C.** Live Ca²^+^ imaging, using GCaMP6s, of TDN and A19c responses to optogenetic activation of Basin-2. **C1**. imaging of TDN response (n=2 animals). **C2**. imaging of A19c response (n=3 animals). Light lasted for 5s. **D.**Live Ca²^+^ imaging, using GCaMP6s, of A19c and TDN response to mechanical stimulation (piezoelectric-delivered vibrations) known to recruit chordotonal mechanosensory neurons. **D1**. imaging of TDN response (n=10 animals). **D2**. imaging of A19c response (n=8 animals). Stimulations lasted for 5s. **E.** Imaging A19c and TDN neurons upon their optogenetic activation and simultaneous presentation of mechanical stimulation **E1.** imaging of TDN responses (n=8 animals) **E2.** Imaging of A19c responses (n=8 animals).

We further determined the neurotransmitter identities of the A19c and TDN using immunolabeling. Both TDN and A19c are glutamatergic (Supplementary fig. 7B), and therefore could be inhibitory^41^. TDN is also cholinergic, and it could also be excitatory (Supplementary fig. 7A). Both neuron types were negative for the GABA neurotransmitter (Supplementary fig. 7C).

### Thoracic and abdominal neurons in the R11A07 line have distinct functions in defensive behaviors

We tested the connections between mechanosensory neurons and TDN and Basin-2 neurons and A19c functionally using calcium imaging in intact larvae. We first investigated the functional connectivity between Basin-2 and A19c by optogenetically activating Basin-2 using CsChrimson and imaging the calcium responses in A19c and TDN neurons with GCAMP6s. In line with synaptic connectivity data from EM, optogenetic activation of Basin-2 induced calcium response in A19c but not in TDN (Fig. 6C). We then monitored the response of the two neurons to a mechanical stimulus. Strong responses of the thoracic neurons to the stimulation were observed, but no response from the abdominal neuron (Fig. 6D). These results are consistent with the mechanosensory > TDN connections we observed in the EM images. No responses to the mechanical stimulus were detected in the A19c neurons, which is consistent with the fact that this neuron receives no direct mechanosensory inputs. However, A19c receives inputs from Basin-2 that themselves receive mechanosensory inputs^17,18^. The lack of responses in A19c could be explained by the inhibition of A19c by neurons receiving mechanosensory inputs. Behavior data (Fig. 3) point to the competition between the Escape responses evoked by optogenetic activation of the R11A07 neurons and the mechanosensory induced responses. The higher the mechanical stimulation relative to the level of optogenetic activation, the stronger the inhibition of the escape responses (Fig. 3). Since the TDN is activated by mechanosensory stimulation and A19c is not, we reasoned that the competition would involve A19c and not TDN. To test this, we monitored responses of TDN and A19c neurons upon simultaneous optogenetic activation using the R11A07 driver and mechanical stimulation at different intensities (Fig. 6E, Supplementary fig. 7D-G). As expected, the calcium responses in TDN resulting from their optogenetic activation were facilitated by mechanosensory stimulations (Fig. 6E1) while A19c activation was inhibited by mechanical stimulation (Fig. 6E2). Similarly to the escape responses in behavioral experiments this inhibition was dependent on the intensity of stimulation (and the level of activation of A19c neurons) (Fig. 6E2, Supplementary fig. 7D-G).

Furthermore, in order to determine which of the two neurons A19c and TDN promote escape, we have used SPARC^42^ to stochastically drive CsChrimson in subsets of the neurons in the GAL4 line (Fig. 7A). Out of 25 larvae tested that showed a response, four performed a Roll, four a C-shape, and five Hunched (Fig. 7B, Supplementary video 1). The remaining larvae performed actions consistent with responses to the light stimulus. We dissected the selected larvae and used immunohistochemistry to reveal the neurons that were activated depending on the exhibited phenotype. In all the larvae that Hunched CsChrimson was expressed in at least one pair of TDN neurons and very few or no abdominal neurons. In the larvae that performed a Roll, no thoracic neuron was labeled while either all or several pairs of A19c neurons were. In the larvae with the C-shape phenotype several pairs of A19c were present and either no or single unpaired TDN neuron could be observed (Fig. 7A, Supplementary atlas, Supplementary table 5). These results point to a role of TDN neurons in triggering Hunch and A19c neurons triggering Escape behaviors (Fig. 7C). Altogether these results show that the A19c, and not TDN, promotes escape actions in a context dependent manner. In addition, A19c could inhibit Hunching, while TDN could promote it.

**Figure 7.**
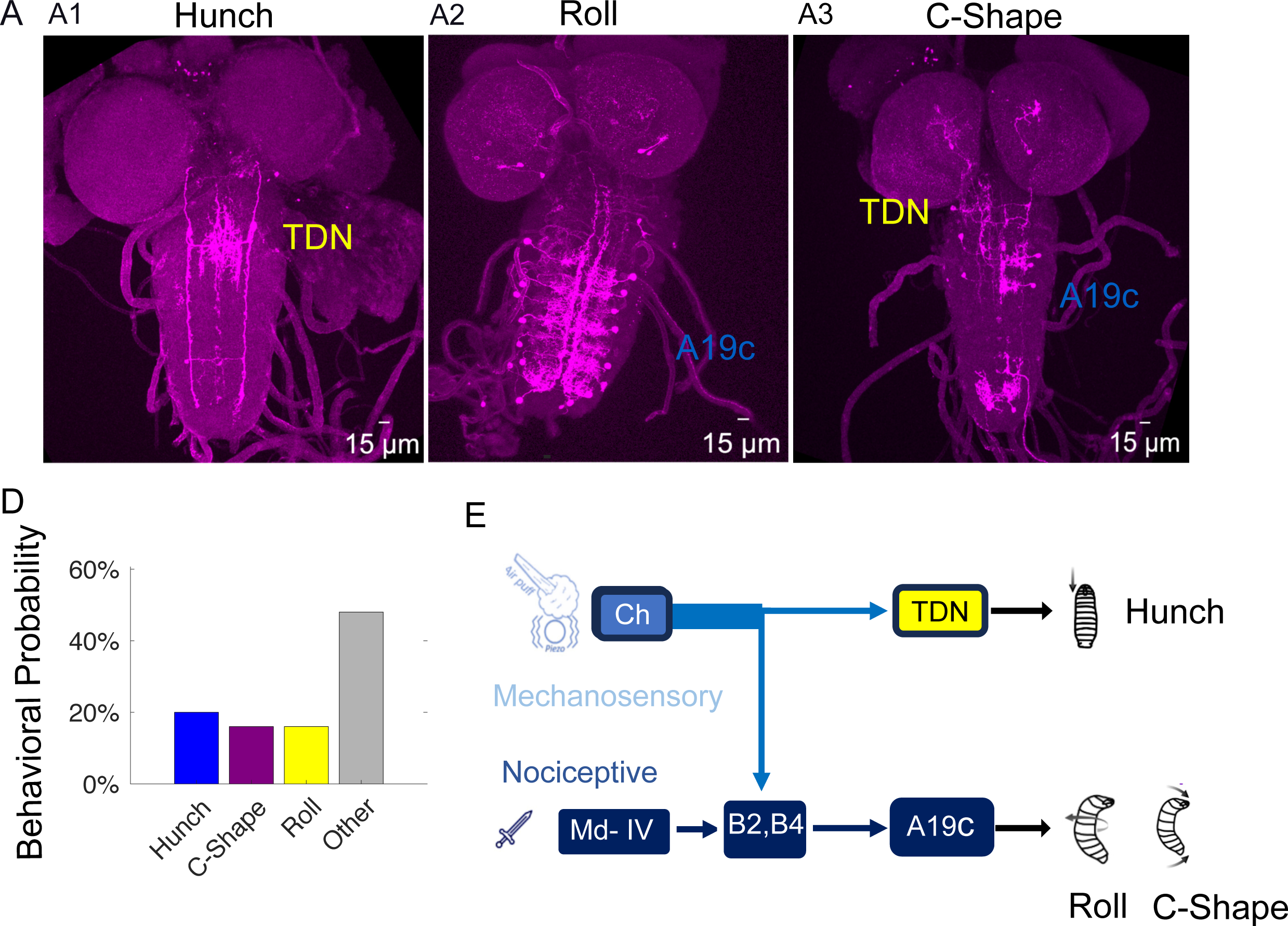
A19c is sufficient to induce rolling. **A.**Example of expression profile of CNS for larvae that **A1.** Hunched, **A2**. Rolled or **A3.** Performed a C-shape **B**. behavioral probabilities of larval responses to optogenetic activation **C.** Schematic showing that TDN promotes Hunching while A19c promotes escape behaviors (y,w;{nSyb-phiC31} attp5;11A07- Gal4::UAS-SPARC2-I-CsChrimsontdtomato} CR-P40 and y,w;{nSyb-phiC31}attp5;11A07- Gal4::UAS-SPARC2-S-CsChrimsontdtomato}CR-P40, **n**=25 animals)

### Selection of avoidance actions is gated downstream of Basin neurons in a context dependent way

Projection neurons Basin-2 and Basin-4 (Fig. 1) were previously shown to inhibit the Hunch response to air puff^17^. Since we found that A19c received significant input from Basins 2 and ™4 (Fig. 1 B-D), was activated by Basin-2 (Fig. 6 C) and that it could also inhibit Hunch like its presynaptic partners (as silencing the neurons in the R11A07 line resulted in more Hunches and its activation suppressed Hunching) (Fig. 1F, Supplementary fig. 2), we hypothesized that Basin-2 and/or Basin-4 could inhibit Hunch through A19c. To test this hypothesis, we optogenetically activated Basin neurons using the LexA drivers (L38H09 for Basin-2 and L72F11 for all Basins) while inactivating A19c with TNT using the R11A07 driver (Fig. 8). As expected, upon Basin-2 or all Basins optogenetic activation during air puff responses, the Hunch was inhibited compared to the respective no-retinal control (Fig. 8A, Supplementary fig. 9 B3,4,C3,4, Supplementary fig. 10 B3, 4, C3, 4). The inhibition was stronger when all Basins were activated than when Basin-2 alone was activated (Fig. 8.A. Supplementary fig. 9, 10B3,4, C3,4). This may be because the combined activation of Basin-2 and ™4, has a greater inhibitory effect than the activation of the Basin-2 neuron alone. The inactivation of A19c and TDN partially rescued the Hunching probability upon activation of all Basins, but not upon activation of Basin-2 alone. This result suggests that the inhibition of Hunch by Basins could be at least partly mediated by their postsynaptic partner A19c. It is worth noting that the Hunching probabilities of the no-retinal control are lower than the Hunching probability of the genetic control (without the LexA) (Supplementary fig. 9A, 10A).

**Figure 8.**
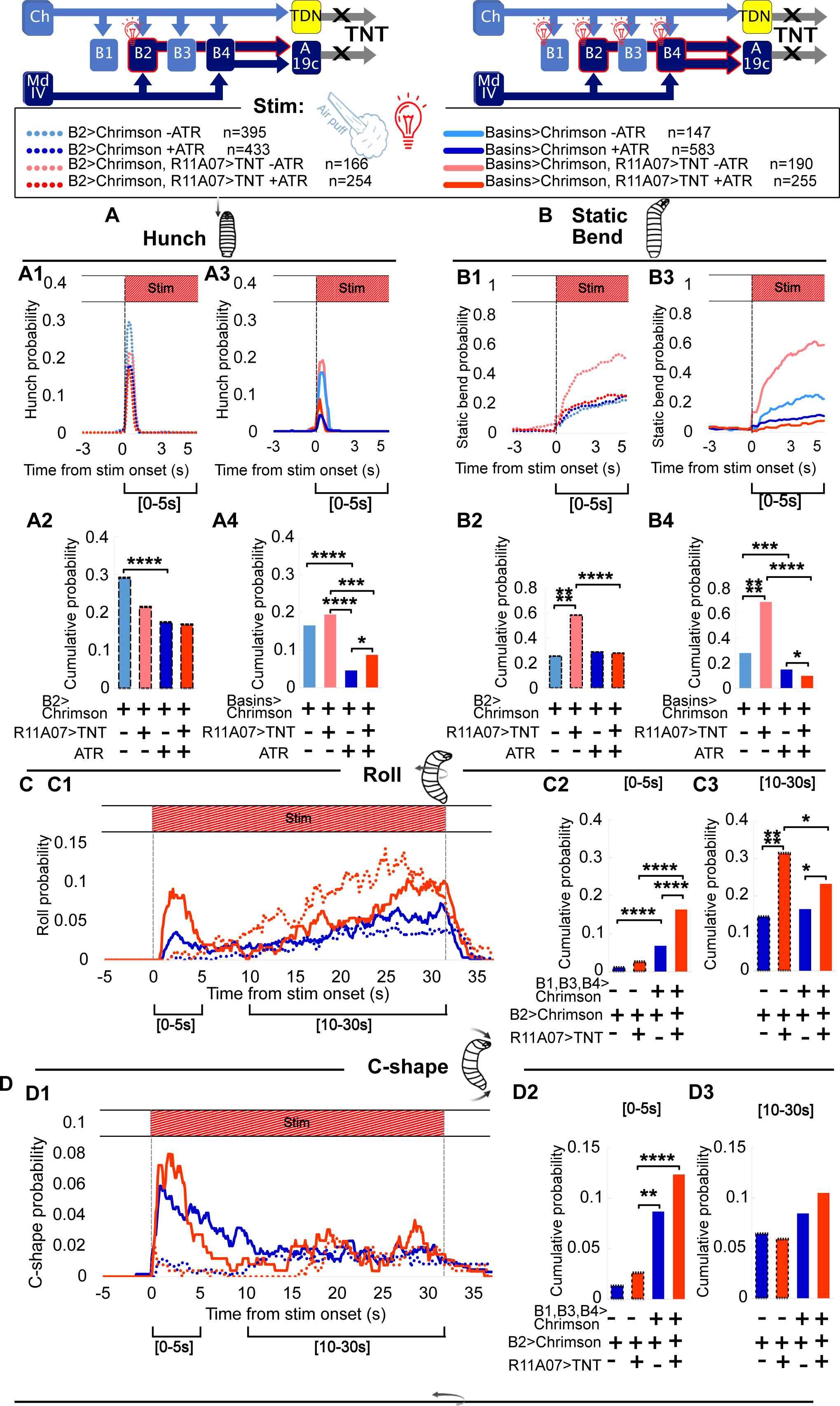
Effect of Inactivating neurons in the R11A07 line on the behaviors triggered by optogenetic activation of Basins. **A.** Hunching Response to air puff and optogenetic activation of Basins, with and without R11A07 inactivation. **A1.** Hunching probability over time, in response to air puff and Optogenetic activation of Basin-2 (L38H09>CsChrimson, dark blue, dotted n=433), and with R11A07 inactivated (L38H09>CsChrimson, R11A07>TNT red, dotted, n=254). Respective no All-Trans-Retinal (ATR) controls: L38H09>CsChrimson, light blue, n=395 and L38H09>CsChrimson, R11A07>TNT, light red, n=166. **A2.** Hunching probability cumulated over the first 5 seconds after onset of air puff and light, same color code as in **A1. A3.** Hunching probability over time, in response to air puff and Optogenetic activation of all Basins (L72F11>CsChrimson) (dark blue, n=583), with R11A07 inactivated L72F11>CsChrimson, R11A07>TNT (red, n=255). Respective no All-Trans-Retinal (ATR) controls: L72F11>CsChrimson, pale blue, n=147 and L72F11>CsChrimson, R11A07>TNT pale red,n=190. **A4.** Hunching probability cumulated over the first 5 seconds after onset of air puff and light, same color code as in **A3 B.** Static-Bend response to air puff and optogenetic activation of Basins, with and without R11A07 inactivation, color code and animal numbers as in **A B1.** Static-Bend probability over time, in response to air puff and Optogenetic activation of Basin-2, with and without R11A07 inactivation. **B2.** Static-Bend probability cumulated over the first 5 seconds after onset of air puff and light **B3.** Static bend probability over time, in response to air puff and Optogenetic activation of all Basins (L72F11) with or without inactivation of R11A07 neurons **B4.** Static Bend probability cumulated over the first 5 seconds after onset of air puff and light **C.** Rolling probability in response to air puff and simultaneous optogenetic activation of Basins, with and without inactivating R11A07, color code and animal numbers as in **A. C1.** probability over time **C2.** Rolling probability cumulated over the first 5 seconds after stim onset. **C3.** Rolling probability cumulated over [10-30s] within stim. **D.** C-shape probability in response to air puff and simultaneous optogenetic activation of Basins, with and without inactivating R11A07. Color code and animal numbers as in **A. D1.** probability over time **D2.** C-shape probability cumulated over the first 5 seconds after stim onset.

The inactivation of neurons in the R11A07 line results in an increase in Static Bending (Fig. 8B, Supplementary fig. 9B3,B4,C3,C4, Supplementary fig. 10B3,B4,C3,C4). Activating Basins decreased the Static Bending probabilities whether the TNT was expressed in R11A07 neurons or not (Fig.8 B3,4, Supplementary fig. 10B,C). This suggests a parallel pathway of Static Bending inhibition by Basins that is not mediated by A19c.

As previously said, the optogenetic activation of Basins was shown to trigger Rolling^18^. Having found that Basins’ inhibition of Hunching could be mediated by A19c, we sought to test whether the same neuron also mediated the Escape sequence. Surprisingly, we found that expressing TNT in the R11A07 neurons dramatically increased Rolling and C-shapes upon Basins activation, both when delivering air puff combined with light (optogenetic activation) (Fig. 8C, D) and light alone (Fig. 8E,F, Supplementary fig. 9, 10 B1,B2,C1,C2). This result suggests that the neurons in the R11A07 can not only trigger C-shapes and Rolling, as shown earlier (Fig. 1-3) but can also inhibit it. Moreover, it seems that the Basin neurons-mediated C-shapes and Rolling may be inhibited by the R11A07 neurons and potentially by their downstream partner A19c.

Interestingly, we found very few Rolls upon optogenetic activation of Basin-2 and all Basins with light alone (Fig. 8D). Rolling was however present upon optogenetic activation and simultaneous air puff stimulation (Fig. 8C). This is consistent with previous work showing that multisensory integration (combined noxious stimulation and vibration) enhances the escape sequence^18^. Thus, similarly to vibration, air puff could enhance the Rolling response upon Basin activation. Optogenetic activation of all Basins and Basin-2 triggered C-shapes and an increase in Head casting (Supplementary fig. 9B,C, 10B,C). Examining the head and tail speed of Head Cast events revealed ‘Tail Casts’ especially in larvae with optogenetically activated Basin-2 neurons (Supplementary fig. 11A) and faster Head Cast. Examining the Crawling speed prior to and upon optogenetic activation revealed Fast Crawls triggered by Basin activation (Supplementary fig. 12). The speed increased upon expressing TNT in R11A07 neurons. As mentioned previously, the Fast Crawls were also induced by optogenetic activation of R11A07 neurons (Supplementary fig. 4). Thus, A19c could inhibit both Rolling and Fast Crawling upon Basin optogenetic activation.

A potential explanation for the absence of Rolling upon optogenetic activation of Basins in our hand compared to previous work could be the use of different drivers and /or different effectors (L72F11, L38H09 and Chrimson in this study and TrpA and R72F11 in Ohyama *et al.*, 2015). In addition, the L38H09 (LexA) labels the Basin-2 neurons stochastically in different segments which is likely to have an effect on the strength and the type of the phenotype observed (for example the Tail Casts).

Altogether we found that neurons in the R11A07 line could mediate the inhibition of Hunch by Basins and they also inhibit C-shapes, Rolls and Fast Crawls triggered by Basin optogenetic activation.

### TDN and A19c neurons connect to the pre-motor and motor layers through distinct pathways

Calcium imaging and SPARC experiment showed that TDN and A19c are differentially involved in Startle and Escape actions (A19c promotes C-shapes and Rolls, while TDN promotes Hunching) (Fig. 6 and 7). In addition, both R11A07 neurons’ optogenetic activation and silencing upon Basin activation induced Rolling also suggesting opposing roles of these neurons in Rolls. To investigate this, we analyzed the postsynaptic connectivity of TDN (Fig. 9, Supplementary fig. 13, Supplementary table 4) and A19c (Fig. 7, Supplementary fig. 13, Supplement table 4) in the EM images. This analysis revealed that TDN is directly connected to motor neurons, whereas A19c is not (Fig. 9, Supplementary table 4). Since TDN receives direct sensory input from multiple segments (Fig. 6, Supplementary table 1), this suggests that TDN could trigger behaviors based on direct mechanosensory inputs. The main direct partners downstream of TDN are premotor neurons T19v and A19d. The premotor neuron T19v, receives inputs from multiple neurons previously described as triggering Rolling, namely Wave^43^ and Basin-2, ™4 and pre-goro neurons^18^. Moreover, T19v is the main premotor target of Wave and Basins-2 and ™4. Thus, TDN could be inhibiting Rolling by influencing T19v.

**Figure 9.**
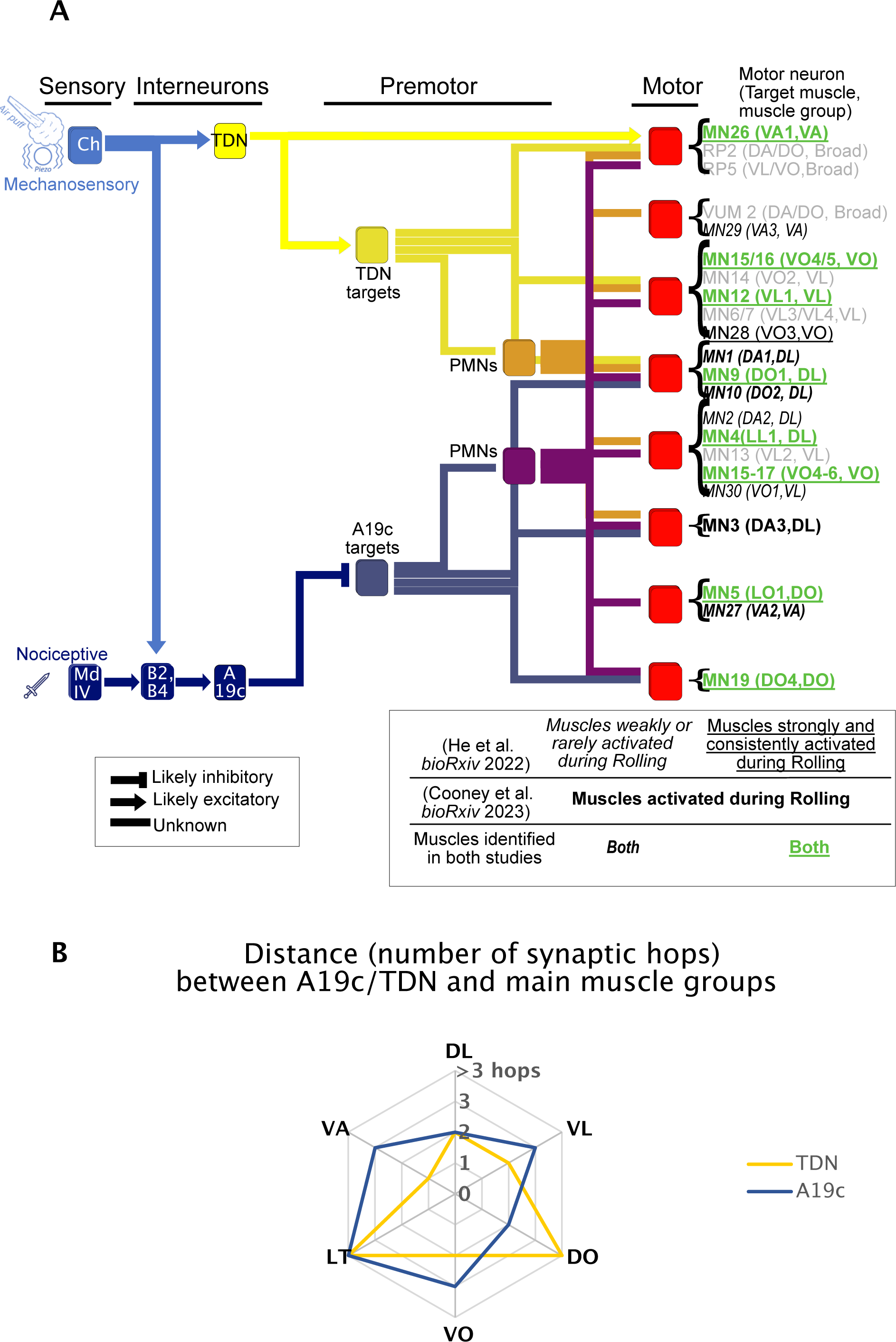
TDN and A19c neurons connect to the motor side through distinct pathways. A. Schematic summarizing TDN and A19c connectivity, and their respective synaptic distance to specific motor neurons. This schematic is a simplified representation of the connectivity shown in fig.7 supplement 1. Postsynaptic partners of TDN were grouped based on whether they were direct (yellow) or indirect (orange) targets of TDN. Similarly, A19c direct (blue) and indirect (purple) targets were regrouped. TDN and A19c contact, directly and indirectly, 22 motor neurons (MNs) located in segment A1. Note that, just as in Figure 7 supplement 1, the motor neurons from other segments were not taken into account. The 22 A1 MNs contacted by TDN and/or A19c were placed into 8 groups (red), depending on their synaptic distance to TDN and A19c. Importantly, among these 22 MNs are MNs targeting muscles identified as recruited during Rolling events, in two recent studies (He et al., BioRxiv 2022; Cooney et al., BioRxiv 2023). These “Rolling MNs” are coloured and highlighted accordingly. In gray are MNs controlling muscles found not to be activated during Rolling by either study, as well as MNs innervating broad sets of muscles (RP2, RP5 and VUM2 MNs). **B.** Radar chart summarizing synaptic distance of TDN and A19c from the main muscle groups.

The main direct partners downstream of A19c synapses are premotor neurons Marathon and A08e1 (Fig. 9). Interestingly, the premotor neuron A08e1, belongs to a group of late-born ELs interneurons, previously described in Heckscher *et al*., 2015 as involved in left-right symmetric muscle contraction^24^.

The main direct targets of TDN were either premotor (three neurons, 29 synapses) or motor (two neurons, 16 synapses), with just one main target being an interneuron not connected to motor neurons, receiving 18 synapses from TDN (Fig. 9A, Supplementary Table 4). The main direct targets of A19c were interneurons not directly connected to motor and premotor neurons (Fig. 9, Supplementary fig. 13, Supplementary Table 4). These results suggest that TDN is more strongly connected to the motor side than A19c, both directly and indirectly.

In terms of muscle targets, the motor neurons targeted by the TDN (and its partners A19d and T19v) innervate broad muscles groups both dorsally and ventrally (RP2 and RP5 muscles respectively) as well as VL, VO (i.e. VO 15, 16) and DL (9, 10) muscles. The A19c premotor neuron targets connect to MNs innervating primarily DL (1, 9, 10) and DO (5,11,19) muscles. These DOs are not targeted by TDN partners. Two recent studies have identified muscle pattern activity during escape Rolling^40,44^. Silencing MN neurons innervating VL and DO by Cooney *et al* blocked bending and Rolling which is consistent with a role of ventral longitudinal muscles in Rolling. In addition, Cooney *et al*., showed that silencing MN innervating DL reduced Rolling. He *et al* identified DL9 as one of the 11 muscles robustly active during Rolling while DL1 and DL10 are less robustly activated. He *et al*. described a total of 11 muscles belonging to the VL, VO, DLA and VA groups that were active during Rolling. Out of these 11 muscles 6 (DL 9, 4, VL 12, VO 15, 16 and 28) were targeted by MNs receiving indirect inputs from both A19c and TDN, while 3 (DO 5, 11, 19) were only targeted by neurons receiving indirect input from A19c.

Together, these connectivity analyses reveal distinct pathways for TDN and A19c to motor neurons, with TDN being closer to the motor side than A19c and TDN recruiting motor neurons more broadly than does the A19c. Interestingly, both A19c and TDN indirectly connect to DL and VO muscles putatively involved in Rolling, although through distinct premotor neurons. In addition, A19c indirectly connects to DO-innervating MN involved in Rolling.

## Discussion

Using machine-learning-based algorithms for behavioral classification combined with targeted neuronal manipulations, EM connectomics, and Calcium-imaging, we characterize the competition between static and active defensive behaviors at the behavioral and neural circuit levels. We identify second-order interneurons that could be differentially involved in Startle (Static) and Escape (Active) behaviors during responses to an aversive mechanical stimulus, the air puff. Based on connectomics analysis, we further identify a descending neuron that could be involved in modulating the escape sequences based on mechanosensory information and inputs from the brain.

### Pathways for static and active defensive behaviors

To evade potential dangers, animals employ diverse defensive strategies from static protective actions to vigorous and fast active forms of escape behavior^5^. The selection of behavior that will be performed will depend on the type of danger as well as the specific context or animal’s behavioral or internal state. Neuronal circuitry that ensures the selection between these different options is not extensively characterized, but competition between freezing and flight has been proposed to involve reciprocal inhibitory connections between two populations of inhibitory neurons for example^13^. Previous work has identified a similar motif with synaptic and cellular resolution and mapped the circuit that underlies the competition between the two most prominent actions that occur in response to the air puff: passive type of response: Hunch, and an active type of response: Bend^17^. In this circuit, the Hunch is encoded by the activity of a projection neuron Basin-1 and in the absence of the activity of Basin-2 at the output of the circuit. The coactivation of both neurons will trigger a Bend (and inhibit the Hunch). Here we identify the next layer in the sensory processing stages composed of two groups of neurons. The early born Even-skipped lateral (EL) neurons that receive primarily Basin-1 (and Basin-3) inputs promote the Hunch, while the A19c neuron that receives primarily Basin-2 and Basin-4 inputs could inhibit the Hunch. Connectivity analyses did not reveal mutual direct or indirect connections between these neurons. By extension, the outcome of the competition at the early processing stages encoded at the level of Basin neurons would be transmitted to the premotor networks to trigger the corresponding behaviors, by inhibiting/activating the appropriate patterns of motor neurons for each behavior. Alternatively, additional sites of competition could exist closer to the motor side.

### Early born EL interneurons could be involved in different types of defensive behaviors induced by mechanical stimulation

EL interneurons were previously shown to be involved in maintaining left-right symmetrical contraction^24^. The early-born EL that receive inputs from chordotonals were shown to respond to vibration^25^ and were proposed to be part of a mechanonociceptive escape circuit: their optogenetic activation triggered Rolling escape. Our results show that early born ELs are both required and facilitate air puff induced Hunching. Thus, early born EL could be involved in different types of defensive behaviors induced by mechanical stimuli. We manipulated the early born ELs using the driver that labels the five different early born neurons. Two of these receive mechanosensory inputs (from chordotonals and Basin-1 and ™3 neurons) and three receive unknown inputs. It is thus possible that, similarly to Basins, the different types of avoidance actions (active and passive forms of escape) are encoded differently by different types or combinations of different types of early born EL interneurons. In order to investigate this, it would be necessary to have genetic access to individual and /or subsets of early born Els.

### A19c neurons inhibit Hunching and promote Rolling

In this study, we also show that the activation of the two neurons in the R11A07 inhibits Hunching and triggers an escape sequence that, in addition to the previously characterized C-shapes and Rolls^23^ can also be composed of Head-and-Tails, as the first action in a sequence. It remains unclear whether Head-and-Tail is indeed a part of an escape sequence in some natural contexts or whether it occurs only after optogenetic activation of R11A07 where the two neurons A19c and TDN are co-activated.

We show that the relative levels of optogenetic activation of R11A07 neurons and the air puff intensities determine the type of escape sequence triggered as well as the probabilities of Hunch/Static Bend (static protective actions) with respect to the escape sequence. Given that the optogenetic activation of the R11A07 targets A19c downstream of the early processing stage competition site, this could suggest that additional sites of competition between static and dynamic escape actions could exist at the premotor site, where competing neurons are activated by the mechanosensory stimulation and A19c optogenetic activation.

Previous studies have found that Basin-2 and Basin-4 neurons inhibit Hunching during air puff responses^17^ and Basins activation triggers Escape behaviors^18^, In this study, silencing the R11A07 upon all Basin activation did result in a moderate increase in Hunching suggesting that A19c could be involved in the inhibition of Hunching by Basin-2 and ™4 neurons.

Furthermore, stochastically activated subsets of neurons in the R11A07 line confirmed that A19c neurons promoted Rolling while TDN neurons promoted Hunching. Silencing R11A07 did, however, also result in more Rolling upon Basin activation suggesting that R11A07 neurons can also inhibit escape behaviors. Given that we found that during simultaneous R11A07 optogenetic activation and air puff stimulation, air puff inhibited Rolling, while in the case of simultaneous air puff and Basin activation, air puff enhances Rolling, this could suggest that different pathways mediate the Basins-induced escape behaviors and the escape sequence induced by optogenetic activation of R11A07 (Fig. 2). Alternatively, since the R11A07 labels two different neurons: TDN and A19c, these could have different roles in the escape sequence, with one neuron inhibiting Rolling while the other promoting it.

Rolling is thought to be the fastest type of escape action induced by the most noxious stimuli^18,21,22^. Despite the fact that increasing the level of optogenetic activation seems to induce stronger escape behaviors: starting from Head casting at lowest intensities to Rolling at medium and high light intensities, the strongest optogenetic activation of R11A07 neurons triggers a majority of C-shapes and not Rolling. This may be due to the uncoordinated effect of optogenetic activation (simultaneous activation of all segments of both the left and right side) compared to natural stimuli, which may be unilateral and more local, that becomes exacerbated at higher levels of optogenetic activation. In addition, increasing the optogenetic activation could have a differential effect on TDN and A19c neuron activity or possibly on the premotor neurons activated by TDN and A19c, and could thus trigger different behaviors. Also, the TDN is activated by mechanical stimulation while A19c is not. Thus, combining mechanical stimulation with optogenetic activation of R11A07 could result in a relatively higher level of activation of TDN compared to A19c. This higher level of activation of TDN could result in sequences with more Head-and-Tails and fewer Rolls. Since high levels of optogenetic activation resulted also in less Rolling and fewer Fast Crawls (Fig. 3), this could suggest that R11A07 neurons could be gating the escape responses based on the context for example, depending on their level of activation. This dose-dependent modularity of the different actions within the escape sequence is reminiscent of the DnB (Down and Back neuron activation level-dependent C-shape and Rolling shown by Burgos et al^23^. We propose that the sensory context could influence the level of activation of the R11A07 neurons (by activating/inhibiting them) as suggested by the results of calcium imaging experiments, which in turn would gate the behavioral responses.

### Context-dependent competition gates avoidance responses

Context is usually thought of as a combination of internal and additional external information that modifies the stimulus-response relationship^7,20,45^. In this study, we define that the optogenetic activation using R11A07 triggers the escape sequence and the presence or not of air puff represents a different context. However, in nature, the distinction between what is a stimulus and what is context is less obvious and the extra information may be part of the stimulus itself that would induce a change in the behavioral response^7,8,45^. Indeed, it has previously been shown the multisensory integration of mechanical and nociceptive stimuli enhances action-selection^18^. However, regardless of the definition, the investigation of the influence of “context” on stimulus-response pairing sheds light on how the different types and different combinations of information are processed and represented at the neural circuit level. Here we find that the relative levels of activation of the R11A07 neurons (more specifically A19c neurons) and air puff intensities determine the probabilities of the different actions in the action sequence induced by simultaneous air puff stimulation and R11A07 optogenetic activation. Stronger air puff intensities can inhibit the Roll, while high optogenetic activation can inhibit the Hunch. This suggests that A19c and TDN neurons could gate the behavioral selection depending on contextual information towards more active or passive forms of avoidance behaviors for example. That competition is mutual where the nociceptive pathway that typically elicits active and fast forms of escape can inhibit protective Hunching and Static Bending and conversely mechanosensory stimulation modulates the escape sequence and can inhibit escape behaviors in favor of the protective/startle-like actions. A recent study has found that chordotonal activation can gate weak nociceptive inputs by acting on second order interneurons A08n through GABAergic inhibition of nociceptors synaptic outputs^46^. Our results indicate additional sites of cross-modal gating of nociceptive response by mechanosensory stimulation, closer to the motor side. Additionally, they point to mutual competition between mechanical and nociceptive responses.

In addition to the inputs from mechanosensory neurons, TDN neurons receive descending brain inputs through axo-axonic connections, suggesting that it could modulate the motor output based on integrated multisensory information, state, motivational drive and/or experience.

### TDN could modulate the defensive actions based on the context

The activation of the TDN and 19c neurons triggers a Roll while their silencing upon Basin activation reveals that one or both of the neurons could inhibit the Roll and the Fast Crawl (as silencing of these neurons results in more Rolling and Fast Crawling upon Basin activation). Because TDN does not receive input from Basin-2 and generally receives very weak input from Basins, this could suggest that the inhibition of Rolling (and Fast Crawling) upon Basin activation could be mediated by the A19c neuron. However, TDN could inhibit the Roll based on inputs coming from the brain by acting on the Basin downstream directly on the motor side (e.g. T19v neurons). Indeed, TDN responds to mechanical stimulation and promotes Hunching (Fig. 6 and 7) and R11A07 optogenetic activation experiments revealed sensory context dependent mutual competition between Hunching and Escape actions (Fig. 3), it is likely that the inhibition of Roll and fast crawl could be mediated by TDN neurons.

Both neurons connect indirectly to motor neurons innervating muscles shown to be involved in Rolling^40,44^. A19c connects (indirectly) to MN that innervate “Rolling muscles” that are not targeted by TDN partners, suggesting that it could be more significantly involved in Rolling. Cooney *et al*. show that muscle activity during Rolling is synchronous across segments and left-right asymmetric with short periods of left-right hemisegmental symmetry when homologous muscles along the dorsal and ventral midline enter the bend and co-contract. They propose that sequential firing of excitatory and then inhibitory premotor neurons (PMNs) on one side followed by the activation of their counterparts on the other side could underlie the progression of muscle contraction waves around the circumference of the larva. Further functional investigation of the downstream partners of TDN and A19c revealed by the connectome will reveal how each of these neurons influences the patterns of motor activation underlying the Rolling behavior.

## Material and Methods

### Animal rearing and handling

Flies (*Drosophila Melanogaster*) were raised on a standard food medium (ethanol 2 %, methylhydroxybenzoate 0.4 %, yeast 8 %, cornmeal 8 %, and agar 1 %) at 18°C. Third instar larvae were collected as follows: male and female flies from the appropriate genotypes were placed together for mating, then transferred at 25°C for 12-16 h on a petri dish containing a fresh food medium for egg laying. The petri dish was then placed at 25°C for 72 h. Foraging third instar larvae were collected from the food medium by using a denser solution of 20% sucrose, scooped with a paint brush into a sieve and gently and quickly washed with water. Larvae used for optogenetic experiments were raised at 25°C in complete darkness, on standard food supplemented with all-trans retinal at 0.125 mM (reference R240000, Toronto Research Chemicals). The full list of genotypes used can be found in the Supplementary Material 1 Resource table.

### Behavioral tracking

We used an apparatus previously described^17,19,26^. Briefly, the apparatus comprises a video camera (Basler acA2040-90umNIR camera) for monitoring larvae, a ring light illuminator (Cree C503B-RCS-CW0Z0AA1 at 624 nm in the red), a computer and a hardware module for controlling air puff. The arena consisted of a 25625 cm2 of 3% Bacto agar gel (CONDALAB 1804-5) with charcoal (Herboristerie Moderne, 66000 Perpignan) in a plastic dish, and was changed for each experiment. For optogenetic experiments, plates without charcoal were used (so as to be transparent), and larvae were tracked using IR light from below (through the agar). Third instar larvae were collected using 20% sucrose. Collected third instar larvae were washed with water, moderately dried and gently spread on the agar starting from the center of the arena, using a soft-haired brush. Approximately 30–100 larvae were tested simultaneously during each experiment. The temperature of the behavioral room was kept at 25°C. The larvae were tracked using the multi worm tracker (MWT) software (http://sourceforge.net/projects/mwt)^26,47^.

### Air puff stimulation

Air-puff was delivered as described previously^17,19,26^ to the 25625 cm^2^ arena at a pressure of 1.1 MPa through a 3D-printed flare nozzle placed above the arena, with a 16 cm x 0.17 cm opening. The Nozzle was connected through a tubing system to plant supplied compressed air. The strength of the airflow was controlled through a regulator downstream from the air amplifier and turned on and off with a solenoid valve (Parker Skinner 71215SN2GN00) consistent coverage of the arena across experimental days. The air-current relay was triggered through TTL pulses delivered by a Measurement Computing PCI-CTR05 5-channel, counter/timer board at the direction of the MWT. The onset and duration of the stimulus were also controlled through the MWT. Larvae were left to crawl freely on the agar plate for 60 seconds prior to stimulus delivery. Air-puff was delivered at the 60th second and applied for 30 seconds. Larvae were also recorded 60 s after the end of stimulation. Air-flow rates at 12 different positions in the arena were measured with a hot-wire anemometer (PCE-423. pCE instruments) to ensure consistent rates across experimental days. The air puff was triggered through TTL pulses delivered by a Measurement Computing PCI-CTR05 5-channel, counter/timer board at the direction of the MWT. The onset and duration of the stimulus were controlled through an Arduino custom made based interface.

### Optogenetic stimulation

For optogenetic experiments, clear agar plates were used and light was delivered using a custom-made 16×16 617 nm-wavelength LED panel. The arena was also illuminated from below with IR light. so as to use IR from below for tracking and red light for optogenetic stimulation. Light (alone or with air puff) was triggered at the 60th second and lasted for 30 seconds. Light intensity was measured as irradiance (mW/cm²), using a PM16-130 photometer (THORLABS). Irradiance was measured at 12 points across the arena and then averaged. The Light intensity used were 0.1 mW/cm² for low, 0.2 mW/cm² for medium and 0.4 mW/cm² for high intensity.

### Mechano and thermo-nociceptive assays

Experiments were performed with staged 3rd instar larvae as described for mechano-nociception^48^ - and thermo-nociception assays^21,39^. Animals were staged for 6h and allowed to develop for 4 days (96h±3h AEL). For mechano-nociception, larvae that were crawling forward were stimulated on mid-abdominal segments (a3–5) with a 50 mN *von Frey* filament twice within 2s. Each behavioral response was scored as non-nociceptive (no response, Stop, Stop and turn) or nociceptive (C-shape bending, Rolling). Rolling and bending behavior was classified as nociceptive due to their absence in *TrpA1* mutant animals. Stopping, turning or no response were scored as non-nociceptive behaviors^49^. Each genotype was tested multiple times on different days and data from all trials was combined. Statistical significance was calculated using the chi2 test. For thermo-nociception using a local hot probe, a custom-built thermo-couple device was used to keep the applied temperature constantly at 46 °C. Stage and density-controlled 3rd instar larvae (96h±3h AEL) were used and all experiments were performed in a blinded fashion. Larvae that were crawling were touched with the hot probe on mid-abdominal segments (a4–6) until the execution of nociceptive Rolling (up to 10 s). Animals were videotaped and Rolling latencies analyzed in a blinded fashion using ImageJ (NIH, Bethesda). Each genotype was tested multiple times on different days and data from all trials was combined. Statistical significance was calculated using Kruskal-Wallis ANOVA and pairwise comparison with Dunn’s *post hoc* test.

### Behavioral classification and analysis

Behaviors were detected using a custom-made machine learning algorithm that was previously described^5^. Behaviors were defined as mutually exclusive actions. Larvae were tracked using MWT software, all the time series of the contours and the spine of individual larvae are obtained using Choreography. From these times series various features are computed, center of the larva, velocities, etc. All key features are presented in Masson *et al.*, 2020 Behavior classification consists of a hierarchical procedure that were trained separately based on a limited amount of manually annotated data.

Here, we require a more detailed definition of behavior. Hence, we extended the hierarchy with an additional layer to differentiate some bends and Hunches, between different behaviors. Bends were separated into Static Bend, Head Cast (see the description of each behavior below) and Hunches were separated into real Hunches, Head-and-Tail and C-shape.

We take all the Bends and Hunches obtained by the first classification algorithm. In the new algorithm the behaviors will not be subdivide. The whole duration of the behavior will be classified consistently (e.g. if the larva performs a bend for t time, the new algorithm will tag these t times in the same way).

### Action definition

Action definitions from Masson *et al.* 2020 were efficient, **however, we needed to modify some of them.** Hence, our procedure consisted on gathering all Bends and Hunches as defined in Masson *et al*., 2020 and recasting them into the new 5 categories of actions. The last layer was trained to do this reclassification task.

#### Head Cast

Dynamic bends where the head moves laterally from one side to the other. There are two subcategories within Head Cast: one where the head moves strongly but the tail moves at the same speed as the center of mass (slower Head Cast), and another where the tail moves rapidly (fast Head Cast).

#### Static Bend

Low speed turning movement, with minimal head movement, and the angle between the segment between the center of mass and the head and the center of mass and the tail remains constant.

#### Hunch

quick behavior, where the head of the larva retracts, decreasing its length between the center of mass and the head of the larva.

#### C-shape

Position where the larva takes the shape of the letter C *i.e.* the head and the tail bend on the same side.

#### Head-and-Tail

Simultaneous retraction of the head and the tail looking like a Hunch from both sides of the body of the larva.

### New annotated data

We required new annotated data to ensure the classifier matched the phenotype of the larva used in these experiments. The sets of C-shape, Hunch and Head-and-Tail were manually tagged from actions selected using the Masson *et al*, 2020 old behavior classification pipeline. A few numbers of tags are used for the model.

The Head Cast behavior is automatically **tagged** randomly from Bends before the stimulus. Static Bends were actions that occurred only **during** stimulation. The Static Bends is automatically tagged with a threshold on the value of the velocity, if bend **occurs** during n time step, the motion velocity normalized by the length of 50% of ! time step **has** to be under 0.02 s^−2^. We set a low threshold **to avoid influencing** the algorithm with erroneous annotations.

### New features

In order to train the last layer of the pipeline we combined features already evaluated with the previous layer of the classifier **with new features.** All features were averaged **over** the duration of the action, for each type of feature, we extracted the maximum and minimum **values.**

● The three velocities: the head velocity, the motion velocity and the tail velocity normalized by the length of the larva. The motion velocity is the velocity of center of mass return by the MWT software. Head-and-Tail are the terminal point of the spine; The averaging along the spine curve and its derivative, 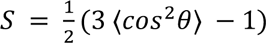 the scalar product between normalized vectors associated to a segment of the spine and the direction of the larva body.
● The shape factor 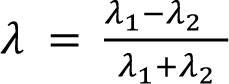 the eigenvalues of the mean covariance matrix of movement which characterizes the shape of the larva and takes value between 0 and 1. All these features are also averaged over 5 time points before and 5 time points after the behavior.

The new features were:

● The ratio between the length of the head-center of mass and the tail-center of mass 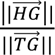 with 1, 2 and 3 respectively coordinate points of the head, the tail and the center of the masse.
● The projection of the Head-and-Tail velocity on the spine of the larva. If we note the velocity vector of the head *HV_h_* with H coordinate point of the head and *V_h_* the coordinate point at the end of the velocity vector, the projection point satisfies the basic relationship:

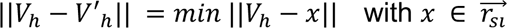

The cosine of the angle between the vector of the head (tail) velocity and the first (last) segment of the larva, 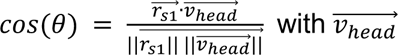 the vector velocity of the head and 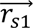 the first vector of the spine.

All features are normalized by the length of the larva to ensure scale-free properties.

### New classifications

We used random forest to perform the classification and divided the process into two steps. A first layer separates original Bends and Hunches into new Hunches, Static Bends and Head Casts. Subsequently, in the second layer, the new Hunches were further categorized into Hunches, Head-and-Tail and C-shape. The performance of the classification is illustrated in Fig. 2A.

We calculated the cumulative probabilities of actions Crawl, Head Cast, Stop, Hunch, Back-up, Roll) (as described in Masson *et al*. 2020) within different time windows after stimulus onset, only in larvae that were tracked at the beginning of the stimulus.

### Statistical analysis

Chi² tests were used for statistical analysis of behavioral probabilities. False rate discovery for applied in case of multiple testing using the Benjamini-Hochberg procedure^50^. When Benjamini-Hochberg procedure was performed, the FDR was reported. The FRD in multiple comparison was not applied in cases when there were a few planned comparisons (Fig. 1 and Fig. 6). In those cases, the p-values were reported.

### Comparison between velocity distribution

We compare some velocity distribution during the Head Cast and the Crawl to identify another type of response to the stimulus. We plot the velocity distribution with a kernel density estimation (using Gaussian kernels). To assess whether two velocity distributions could originate from the same underlying distribution, we employed the two-sample Kolmogorov–Smirnov test.

Two-sample Kolmogorov–Smirnov test is used to compare velocity distribution; it’s a nonparametric hypothesis used to test whether two underlying one-dimensional probability distributions differ. The two distributions come from two different data sets. The Kolmogorov– Smirnov statistic is: *D_n,m =_ sup_x_* (|*F_n_(x)* - *G_m_(x)|*)

with *F_n_(x)* and *G_m_(x)* are the empirical distribution functions of the first and the second sample respectively, and sup is the supremum function. The null hypothesis assumes that the data from the first sample and the second sample are from the same continuous distribution. The p-value is computed from the value of *D_n,m_*.

### Calcium-imaging

We used both intact larvae and filet preparation (where the cuticle was left attached to the CNS). For intact larvae imaging, third instar larvae were rinsed in water, and mounted between a 2 cm circular coverslip and a custom-made device that delivers mechanical stimulations in low melting point agarose 4% (melted in phosphate buffer saline), ventral side facing up. Larvae were gently squeezed in this position until agar cooled down, so that the ventral nerve cord could be imaged through the cuticle.

Filet preparation was done as previously described in^17^ (Jovanic et al., 2016). A precise dissection involved a longitudinal incision along the larva’s midline, carefully removing all organs except nerve connections from the CNS to the cuticle (sensory organs and muscles). The cuticle was cautiously stretched and secured at corners for optimal exposure of VNC. The dissection was done in a calcium solution (NaCl 39%, KCl 1.8%, CaCl2 1.1%, MgCl2 2%, TES 57%, Sucrose 61%) to safeguard neuronal function. Also, delicate nerve connections were handled with care to prevent damage.

Mechanical stimulations were generated by a waveform generator (Siglent sdg1032x) connected to a quick-mount extension actuator (Piezo Systems, Inc.), which was embedded in the sylgard-coated recording chamber. The stimulation was set at 1000 Hz, with an intensity of 1 to 20 V applied to the actuator. The amplitude of the acceleration produced by the actuator was measured thanks to a triple axis accelerometer (Sparkfun electronics ADXL313) connected to a RedBoard (Sparkfun electronics) and bound to the Sylgard surface thanks to high vacuum grease. Acceleration was 1.14 m.s^−2^ at 20 V, and 0.61 m.s^−2^ at 10 V. Mechanical stimulations were precisely triggered by the Leica SP8 software thanks to the Leica “Live Data Mode” and to a trigger box branched to the scanning head of the microscope. A typical stimulation experiment consisted in 5 s of recording without stimulation, then 5 s of stimulation, and 5 s of recording in the absence of stimulation.

For optogenetic activation during in vivo imaging, larvae were mounted in the dark, with the least intensity of light possible in the room, to avoid nonspecific activation of the targeted neurons. Optogenetic stimulation of CsChrimson was achieved by a 617-nm wavelength LED (Thorlabs, M617F2), controlled by a LED driver (Thorlabs, LED1B) connected to the waveform generator, and conveyed through a Ø 400 µm Core Patch Cable (Thorlabs) to the imaging field. Optogenetics stimulations were triggered at 50 Hz, 50% and 100% duty during 1 s, concomitantly to mechanical stimulations thanks to the waveform generator. Irradiance was measured at the level of the imaging field at 500 µW using a PM16-130 THORLABS photometer.

Imaging was achieved using a 2-photon scanning Leica SP8 microscope, at 200 Hz, with a resolution of 512 x 190 pixels. At this resolution, the rate of acquisition was 1 frame/s, 2 depending on the experiment. Some experiments were performed at 50 frames/s, with a 256×256 resolution. A set of stimulation of different intensities was repeated the same number of times in each larva (3-8 times depending on the experiment). A resting interval of 60 s was respected in between each stimulation. The orders of the stimulation were randomized in order to exclude potential bias from a specific order of stimulations. The randomisation was made by associating one number to one specific stimulation and creating randomized suites of these numbers using www.dcode.fr. Neuronal processes were imaged in the VNC at the axonal level and fluorescence intensity was measured by manually drawing a region of interest (ROI) in the relevant areas using custom Fiji macros. Data were further analyzed using customized MATLAB scripts. F0 was defined as the mean fluorescence in the ROI during baseline recording, in the absence of mechanical stimulus or optogenetic activation. ΔF/F0 was defined at each time step t in the ROI as: ΔF/F0 = (F(t) - F0)/F0. The means were made using the same number of repetitions per larva, Experiments showing activity before the mechanical stimulations or optogenetics activation, were removed from the analysis

### Immunohistochemistry labeling

To determine the neurotransmitter identity of the neurons, immuno-labeling was performed from the split lines or Gal4 lines crossed to UAS-myr::GFP, or LexA lines crossed to LexAop-myr::GFP. The VNC was dissected out from 3rd instar larvae and fixed with 4% PFA for 45 min at room temperature. After rinsing in PBS, ten minutes permeabilization in PBS-T and two hours blocking in PBS-T-BSA 1%, the CNS preparations were incubated at 4°C (one to three nights) in the first antibodies raised against neurotransmitter and GFP in PBS-T. Then they were incubated at 4°C (one to two nights) in fluorophore-coupled secondary antibodies in PBS-T raised against species of the first antibodies. After rinsing, the preparations were mounted in an anti-bleaching mounting medium (SlowFade Gold, ThermoFisher S36939) under a cover slip. The confocal images were captured with a Leica SP8 confocal laser microscope. Alexa Fluor 488 was excited with a laser light of 488 nm, Cy3 with a laser light of 561 nm, Alexa Fluor 647 with a light of 633 nm wavelength.

### Optogenetic activation using SPARC

For the SPARC (Sparse Predictive Activity through Recombinase Competition) experiment ^42^, y[1] w[*]; P{y[+t7.7] w[+mC]=nSyb-IVS-phiC31}su(Hw)attP5/Cyotb;11A07-Gal4/Tm6TbSb was crossed with TI{20XUAS-SPARC2-I-Syn21-CsChrimson::tdTomato-3.1}CR-P40 and TI{20XUAS-SPARC2-S-Syn21-CsChrimson::tdTomato-3.1}CR-P40. Among the population of third instar larvae, larvae were preselected depending on the exhibited phenotype in response to red light (escape or startle responses). The behavior of selected single larvae upon optogenetic activation was recorded with MWT (as above). For optogenetic activation red light of 617 nm as described was used at 0.4 mW/cm² intensity. For selected larvae dissection of CNS was performed using immunochemistry experiments. The neurons with expressed CsChrimson-TdT tomato were revealed using rabbit anti-DsRed as the primary antibody and anti-rabbit-Cy3 as the secondary antibody.

### EM reconstruction and connectivity analysis

EM reconstruction was performed using a complete CNS serial section transmission EM volume from a 6-hour old *Canton S G1*c 3 c*w1118 [5905]* larva, with a resolution of 3.8nm x 3.8nm x 50nm^18^. We used the web-based software CATMAID^51^ to annotate the synapses onto and from A19c and reconstruct the 3D morphology of neurons presynaptic and postsynaptic to A19c. To do so, we used a previously described methodology^18,36^. Putative TDN neuron was reconstructed up to recognition, a point by which we could confirm that the neuron’s morphology, including position of the soma, dendrites and axon, matched the features of the neuron we observed in light microscopy images. If at least one of hemilateral pairs of neurons received at least 3 synapses from a particular neuron of interest (A19c, TDN, or their partners) we considered them strong downstream partners. For Basin neuron analysis i.e in Supplementary fig. 1, we considered neurons as not reconstructed up to recognition when below 1500 nodes.

## Acknowledgments

We thank the Cardona lab at the MRC LMB, UK, for hosting the EM volume in their CATMAID server which allowed us to reconstruct neurons and synapses. We thank Casey Schneider-Mizell, Laura Herren, Akira Fushiki, Eri Hasegawa, Ingrid Andrade, Aref Arzan Zarin, Avinash Khandelwal, Tara Guillorit and Claire Julliot De La Morandiere for their contribution to the EM reconstruction. We also thank Ellie Heckscher for helpful discussions and comments throughout the project.

## Author contributions

M.L. behavioral experiments and analysis, EM reconstruction and analysis; Writing: results, original draft C.B. behavioral classification and analysis, statistical analysis, writing: methods B.F. Ca-imaging experiments, A.H. Calcium-imaging, SPARC and immunohistochemistry experiments, S.A SPARC and immunohistochemistry experiments, writing: methods, N. D. behavioral experiments and analysis P.S. supervision, writing-edits and revisions J.B. Methodology, supervision, funding acquisition, writing-edits and revision T.J. Conceptualization, analysis, supervision, funding acquisition and project administration; writing: original draft, edits and revision

## Supplementary information

**Supplementary figure 1.**
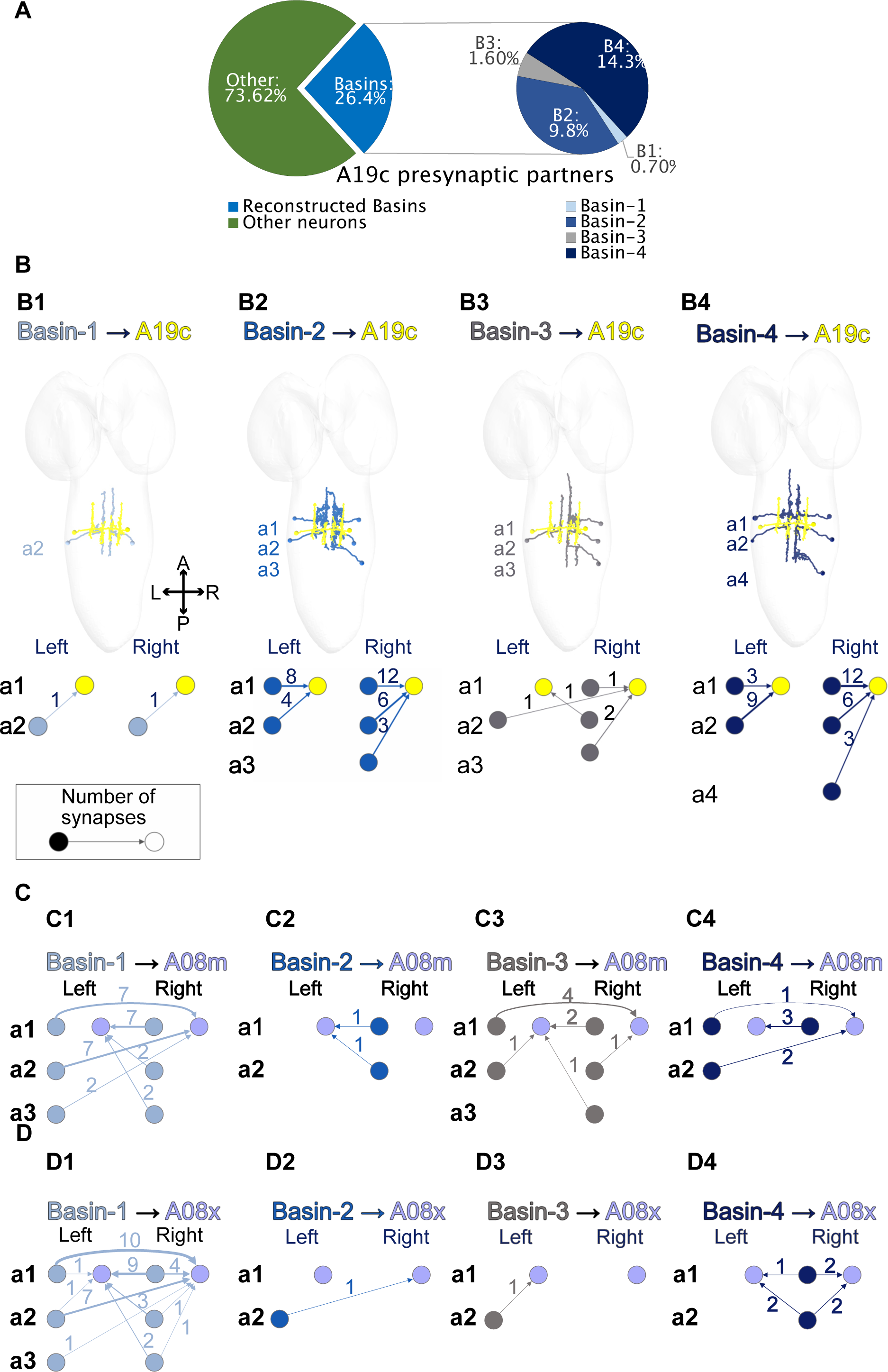
**A.** Distribution of all A19c inputs shown as fractions of inputs of A19c from Basins and non-Basin neurons. **B. B1-B4. Top** EM reconstruction images of Basin neurons in segments and A19c neurons **B1-B4. Bottom** Connectivity of Basin 1-4 - A19c in neuromeres A1-4, left and right. **C. C1-C4.** Connectivity of Basins 1-4 - A08m in neuromeres A1-4, left and right. **D. D1-D4.** Connectivity of Basins 1-4 - A08x in neuromeres A1-4, left and right.

**Supplementary figure 2.**
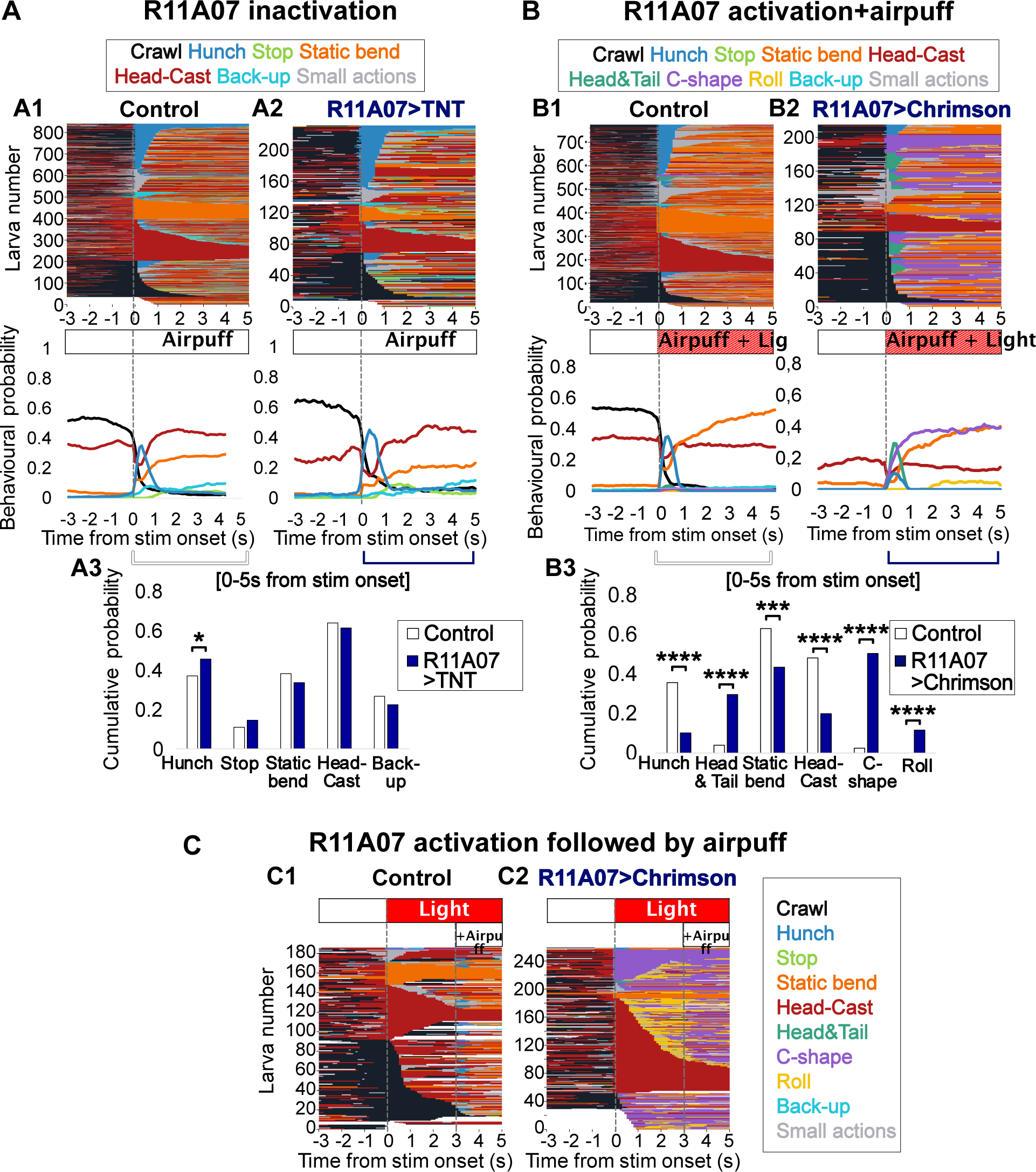
**A.** Larval responses to 4m/s air puff. **A1, A3** control larvae (attP2>TNT, n=818), **A2, A4.** larvae with inactivated R11A07 neuron (R11A07>TNT, n=225). **A1-A2**: ethogram, showing actions over time, with one line corresponding to one individual and each color corresponding to a different action **A3-A4** mean behavioral probability over time. Stim onset at 60 s. **A5.** behavioral probability cumulated over the first five seconds after air puff onset, corresponding to control larvae (white) and larvae with R11A07 inactivated (dark blue). **B.** Larval responses to air puff (4 m/s) and light (0.3mW/cm²). **B1, B3.** control larvae (attP2>CsCrimson, with ATR, n=765, **B2B4.** larvae with R11A07 neurons optogenetically activated (R11A07>CsCrimson, +ATR, n=214). **B1-B2**: ethogram, **B3-B4** mean behavioral probabilities over time. Stim onset at 60th s. **B5,** behavioral probability cumulated over the first five seconds after air puff onset, corresponding to control larvae (white) and larvae with R11A07 optogenetically activated (dark blue). **C.** Larval responses to light (at 60s.) then air puff, delivered 3s after the light activation (at 63 s). **C1** control larvae (attP2>CsChrimson, n=186) **C2.** larvae with R11A07 neurons activated (R11A07>-CsChrimson, +ATR, n=255). For all barplots: *:p<0.05, **:p<0.005, ***:p<0.0005,****:p<0.0001, Chi² test

**Supplementary figure 3.**
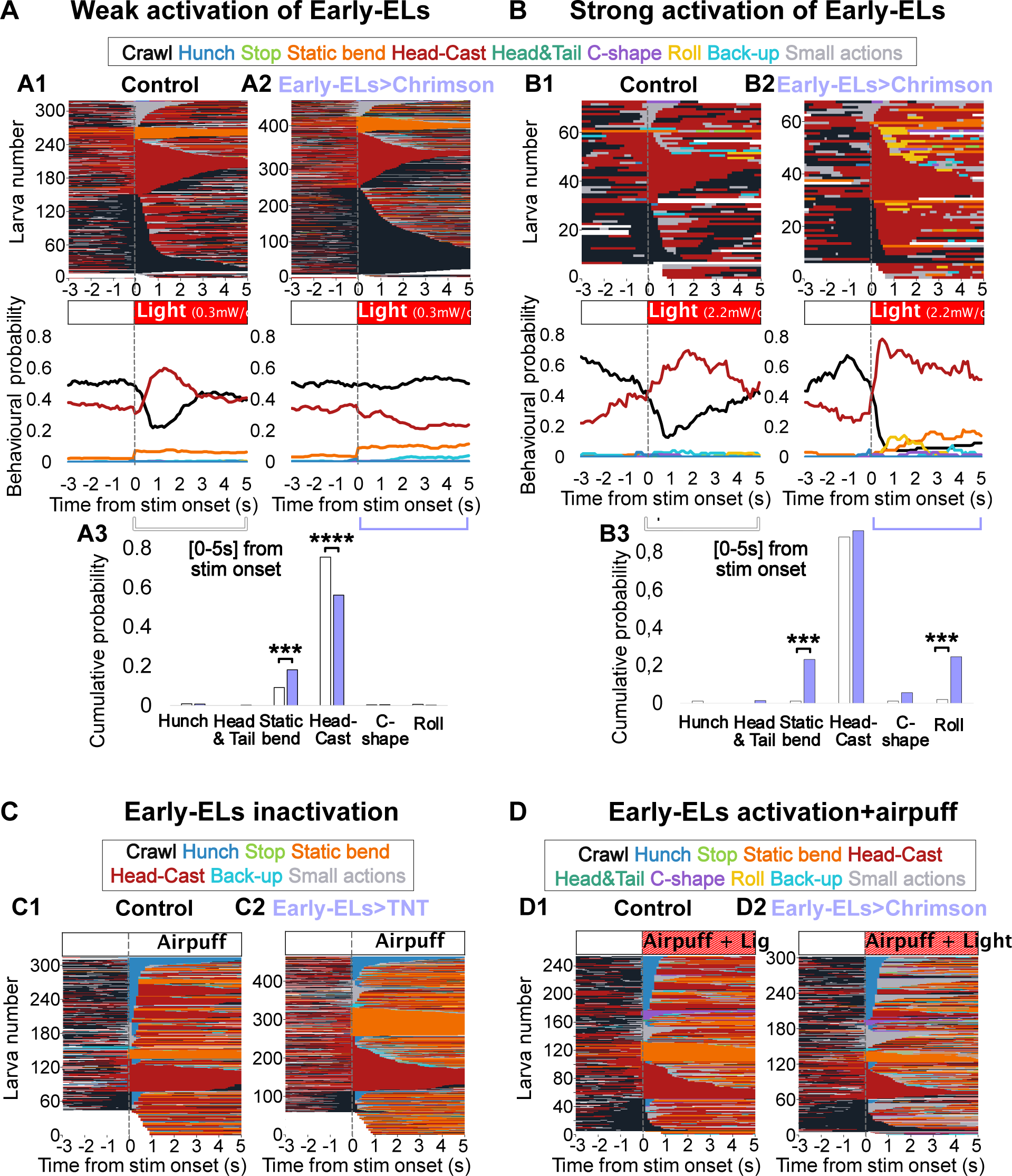
Early-born ELs inactivation and optogenetic activation experiments. **A:** optogenetic activation of early-born eELs (EL-Gal4_R11F02GAL80>CsCrimson) at 0.3mW/cm² irradiance. **A1**: Control (animals reared on food medium without ATR, n=303) Top: ethograms,Bottom: mean behavioral probability over time. **A2**: ELs>CsChrimson (animals reared on food medium supplemented with ATR, n=465). **A3**: behavioral probability cumulated over the first five seconds after light onset**. B:** optogenetic activation of early-born ELs at 2.2mW/cm² irradiance. **B1**: Control (animals reared on food medium without ATR, n=62). **B2**: ELs>CsCrimson (animals reared on food medium supplemented with ATR, n=60). **B3**: behavioral probability cumulated over the first five seconds after light onset**. C.** inactivated early-ELs impacts behavioral responses to strong air puff. **C1**: control (CantonS>TNT,n=254). **C2**: inactivation of early-born ELs (EL-Gal4_R11F02GAL80>TNT, n=420). **D**:early-ELs activation (EL-Gal4_R11F02GAL80>CsChrimson, 0.3mW/cm² irradiance) during strong air puff stimulation 4 m/s **D1**: Control (animals reared on food medium without ATR, n=248). **D2**: eELs>CsCrimson n (animals reared on food medium supplemented with ATR, n=286). Note that behavioral probabilities for **C** and **D** can be found in Figure 1. For all barplots: *:p<0.05, **:p<0.005, ***:p<0.0005, ****:p<0.0001,Chi² test.

**Supplementary figure 4.**
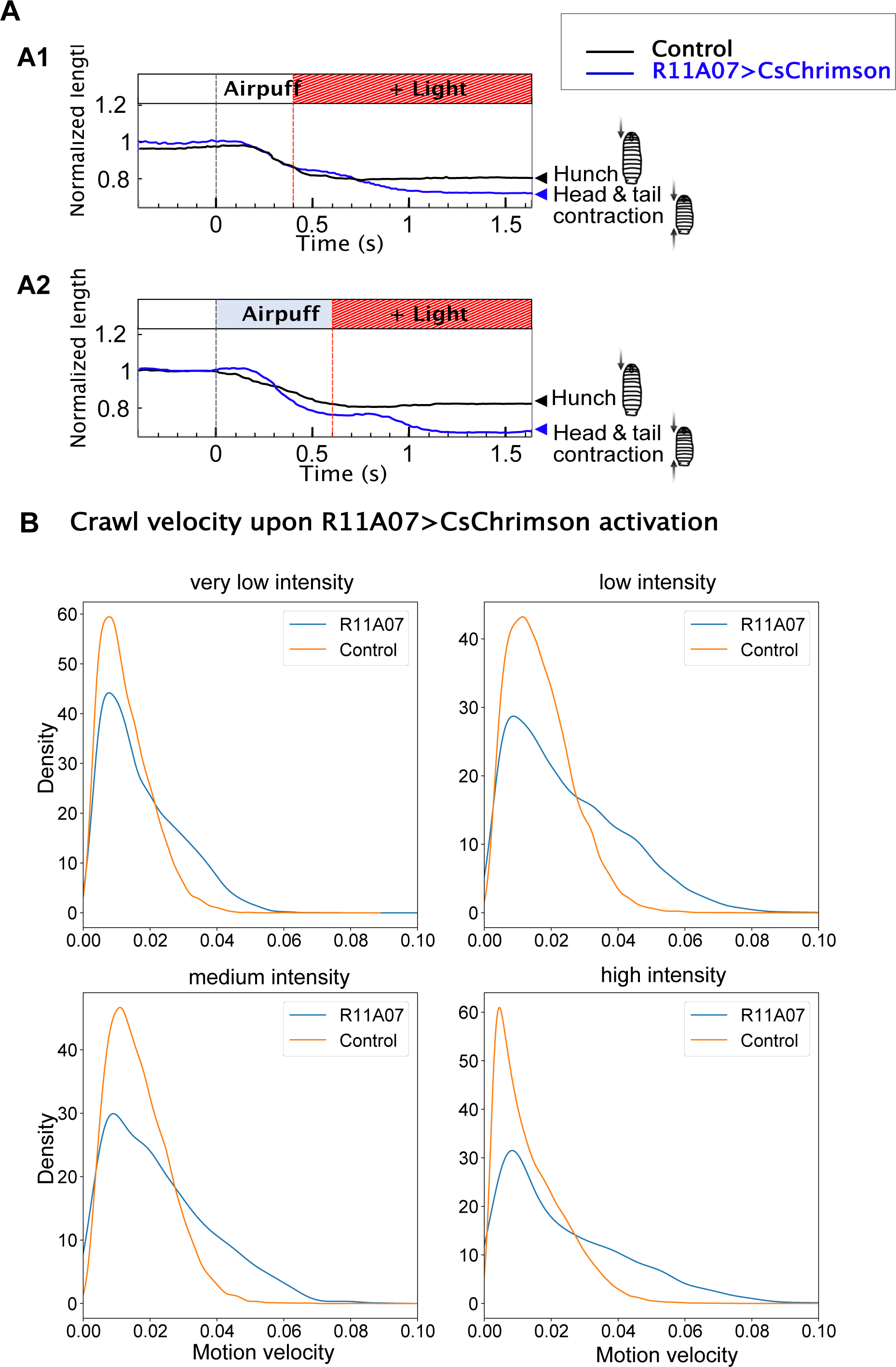
A. Impact of delayed optogenetic activation of neurons labeled by the R11A07 driver on larval length in response to air puff. Control: attP2>CsCrimson, R11A07>CsChrimson larvae were reared on food supplemented with ATR **A1**. Light delivered 0.4s after air puff onset. **A2.**. light delivered 0.6s after air puff onset. **B.** Distribution of motion velocities during Crawls (velocity of the center of mass expressed in normalized body lengths per second (s^−1^)) during the first 10 seconds upon optogenetic activation of R11A07 neurons compared the control. Different intensities of light are shown: very low, low (0.1 mW/cm²), medium (0.2, mW/cm²), strong (0.3, mW/cm²)

**Supplementary figure 5.**
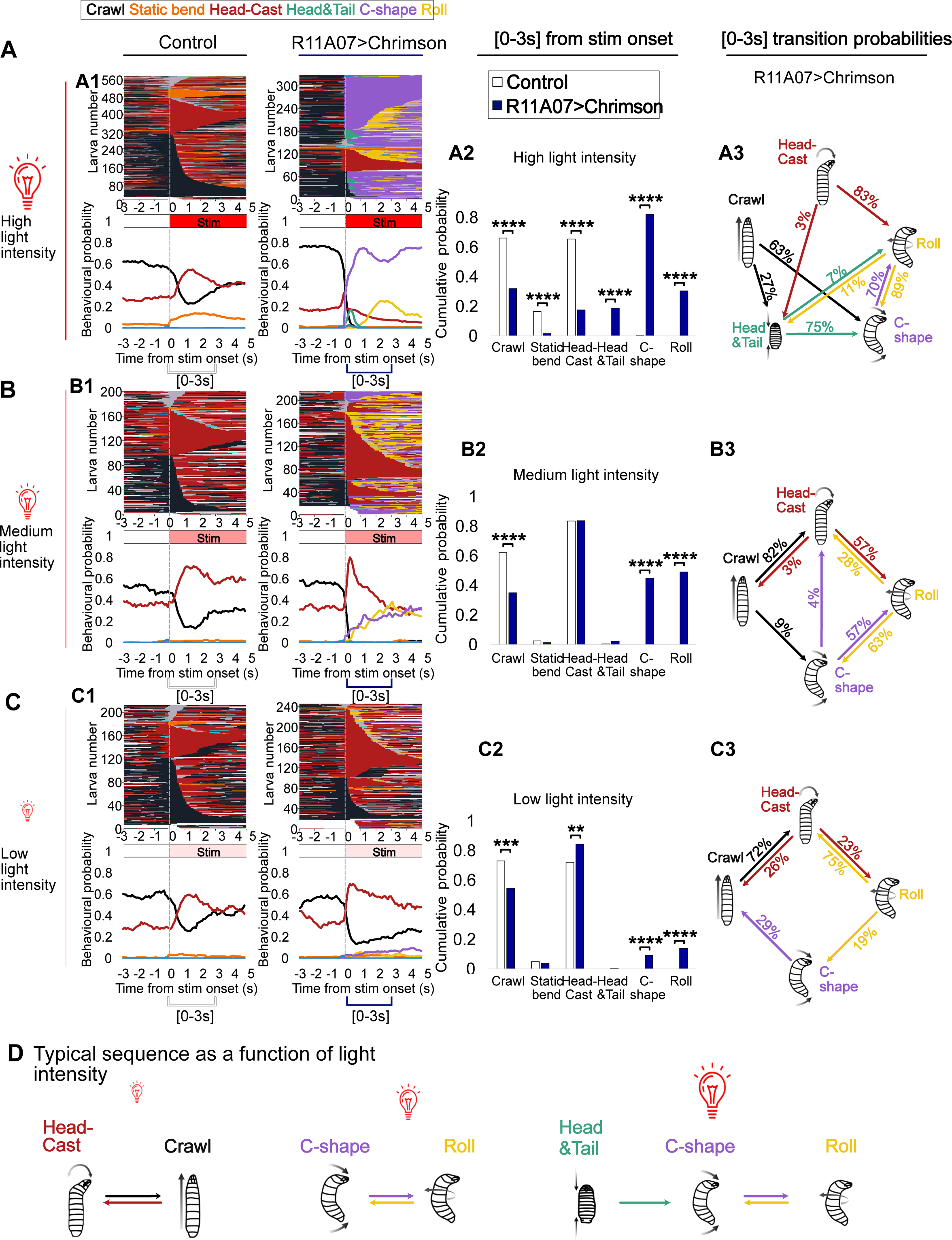
Ethograms, behavioral probabilities over time, cumulative and transition probabilities for different levels of optogenetic activation of R11A07 neurons (R11A07>CsChrimson) and the control (attP2>CsChrimson) **(A.** High, **B** Medium **C** Low light intensity). Cumulative and transition probabilities were computed during the first 3 seconds of light stimulation

**Supplementary figure 6.**
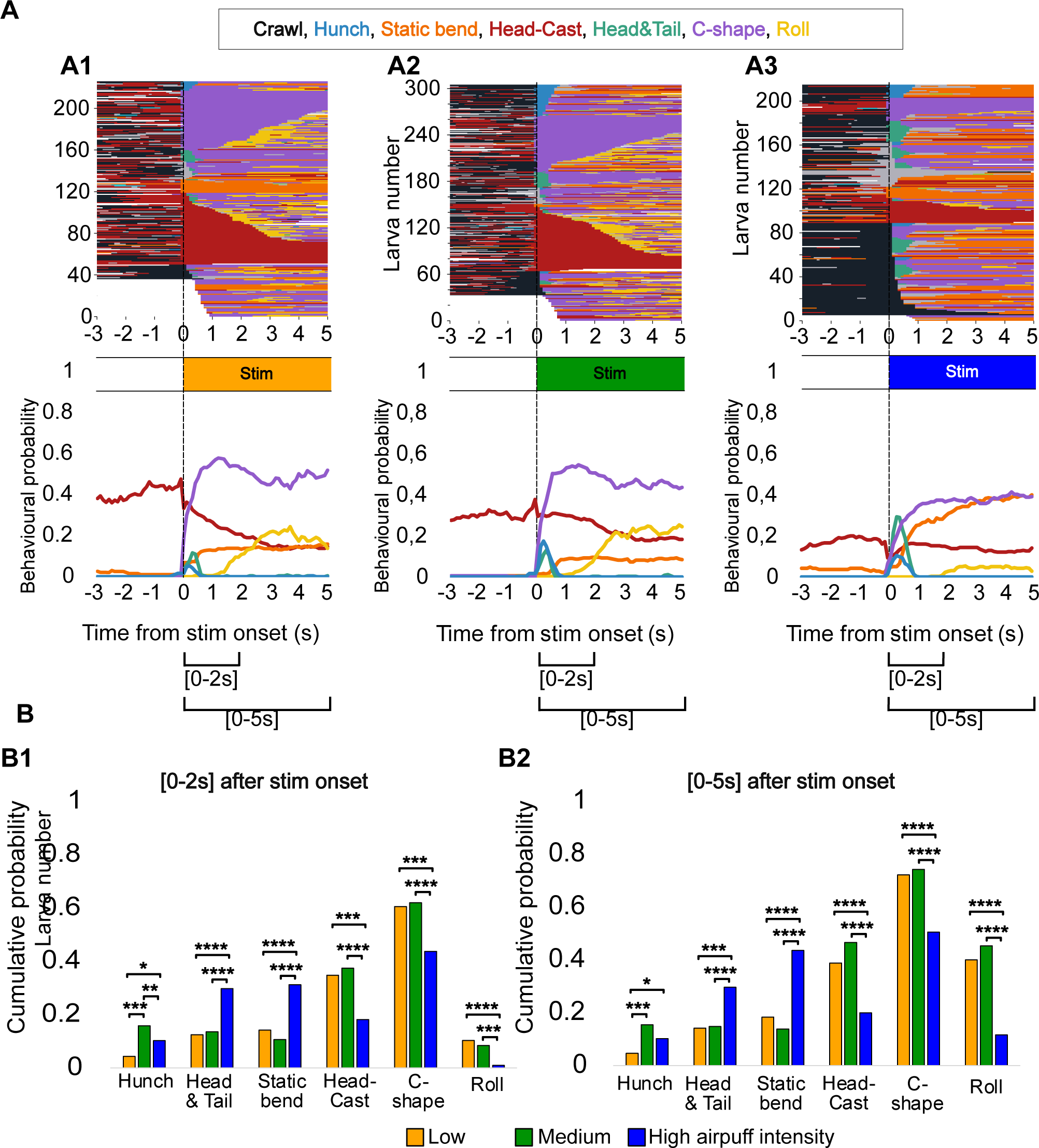
Influence of air puff on escape response induced by R11A07 activation. **A:** behavioral probabilities in response to 0.3mW/cm² light and, **A1.** 2m/s air puff, **A2**. 3m/s air puff, **A3.** 4m/s air puff. **B**. behavioral probability cumulated over: **B1**. the first two seconds after stim onset and **B2**. the first five seconds after stim onset.

**Supplementary figure 7.**
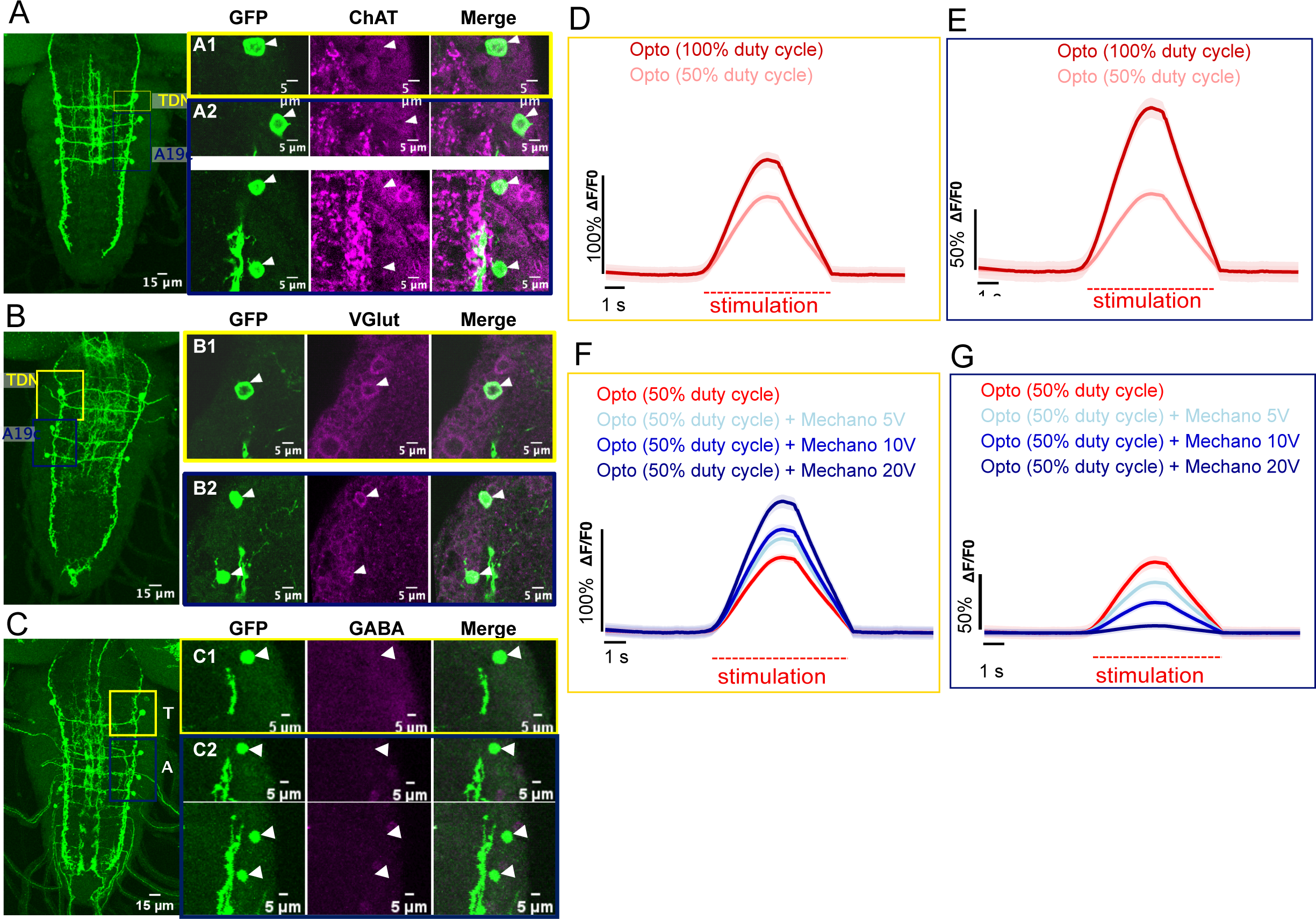
**A-C.** Immunohistochemical labeling of TDN and A19c neurotransmitters. GFP labels the GAL4 expression patterns **A.** 11A07AD; 65E09-DBD>GFP). **B.** 65E09-AD;11A07-DBD>GFP). **C.** R11A07>GFP). Magenta corresponds to neurotransmitter labeling g **A.** Cholinergic (ChAT) **B**. Glutamatergic (VGlut) **C.** GABA **A1, B1, C1**. Thoracic (TDN). **A2, B2, C2**: Abdominal (A19c). **D.** Ca²^+^ imaging, using GCaMP6s, upon optogenetic activation of R11A07 neurons. **D1.** imaging of TDN response (n=8 animals). **D2.** imaging of A19c response (n= 8 animals). Light stimulation lasted for 5s. **E.** Ca²^+^ imaging, using GCaMP6s, upon optogenetic activation of R11A07 neurons and different levels of mechanical stimulation. **E1.** imaging of TDN response (n= 8 animals). **E2.** imaging of A19c response (n= 8 animals). Light stimulation lasted for 5s.

**Supplementary figure 8.**
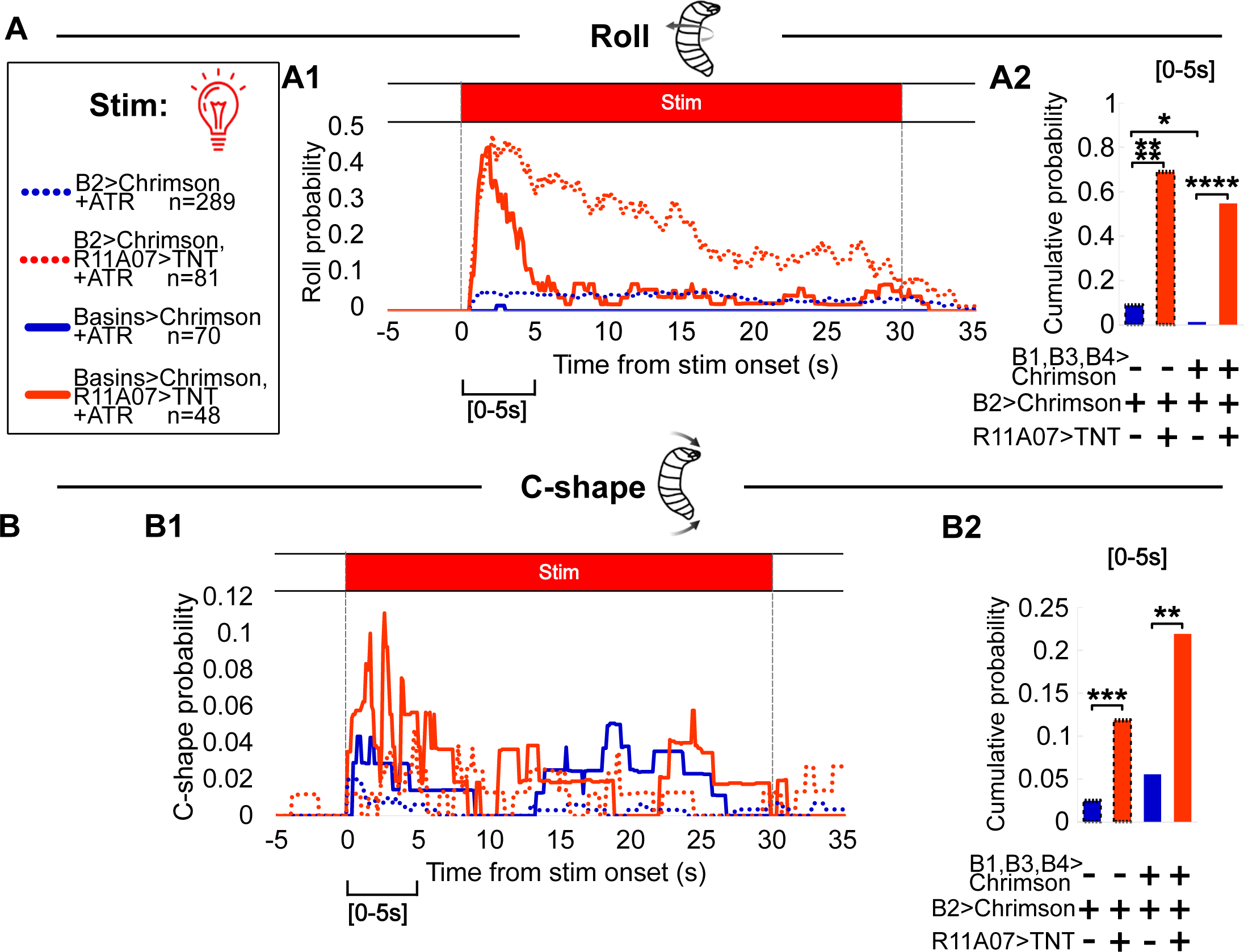
**A.** Rolling probability in response to optogenetic activation of Basins (light alone): Activation of Basin-2 alone (Blue dashed line, n=289), with R11A07 inactivation (Red dashed line, n= 81). Activation of all Basins (blue full line,n=70), with R11A07 inactivation (Red full line, n=48) **A1.** probability over time **A2**. Rolling probability cumulated over the first 5 seconds after stim onset. **B.** C-shape probability in response to optogenetic activation of Basins (light alone). Colors and animal numbers as in A**. A1**. probability over time **A2.** Rolling probability cumulated over the first 5 seconds after stim onset. Light intensity:0.3, mW/cm², airpuff intensity: 4m/s *:p<0.05, **:p<0.005, ***:p<0.0005, ****:p<0.0001, Chi² test

**Supplementary figure 9.**
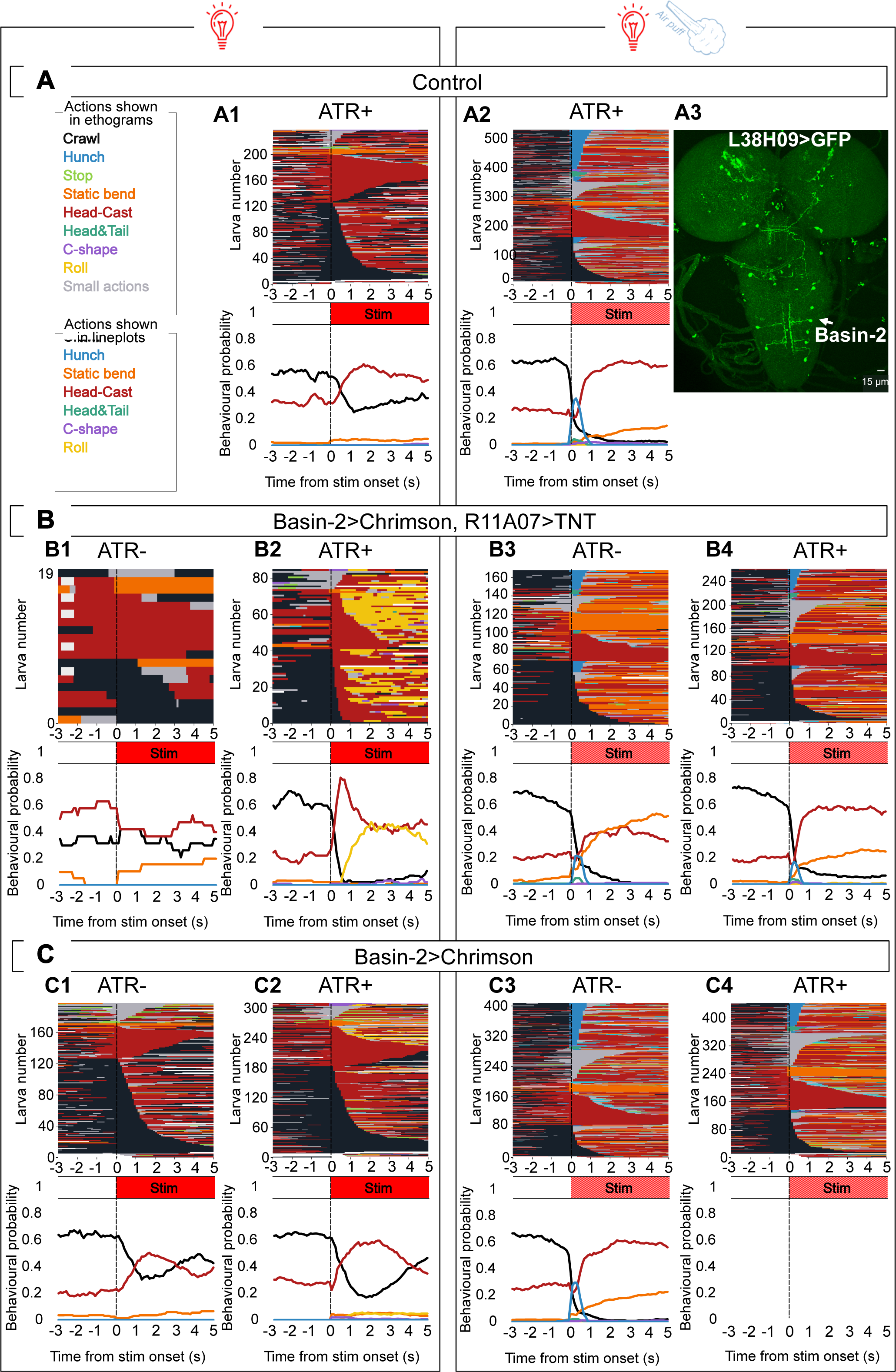
Impact of R11A07 inactivation on the Basin-2 induced responses, Ethogram and behavioral probabilities over time showing all observed actions **A.** Control (attP2-40>CsChrimson, larvae were reared on food medium supplemented with ATR). **A1.** light stimulation (n=199). **A2.** air puff and light stimulation (n=509). **Top**. ethogram, showing behavioral sequences of each individual (one line in the ethogram)as a function of time. Bottom: mean behavioral probability across the population. **A3**. Expression profile of the L38H09 line **B.** optogenetic activation of Basin-2 (L38H09>CsChrimson) with R11A07 inactivated (R11A07>TNT) alone (B1, B2) or combined with air puff stimulation (B3, B4) **B1.** larvae without ATR (n=19), **B2.** with ATR (n=81) during air-puff stimulation: **B3**. without ATR, (n=166), **B4.** with ATR (n=254) **C.** Optogenetic activation of Basin-2 (L38H09>CsChrimson) (C1, C2) or combined with air puff stimulation (C3, C4) **C1.** larvae reared without ATR **C1** (n=170), or with ATR **C2** (n=289). During air-puff stimulation **C3.** without ATR n=395), or **C4.** with ATR (n=433). Light intensity was 0.3mW/cm², air puff intensity 4m/s.

**Supplementary figure 10.**
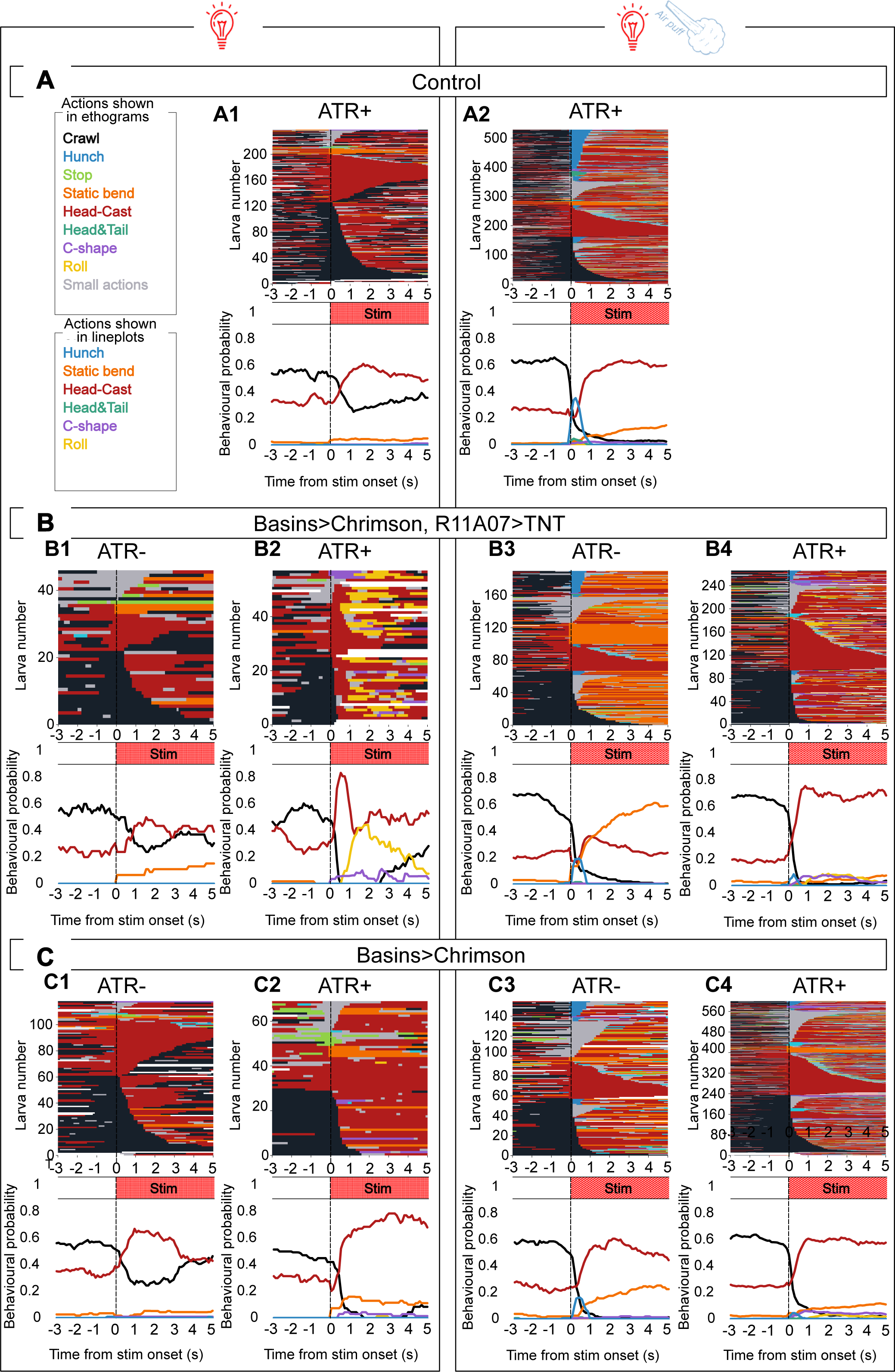
Impact of R11A07 inactivation on the all Basins induced responses, Ethogram and behavioral probabilities over time showing all observed actions **A.** Control (attP2-40>CsChrimson, larvae were reared on food medium supplemented with ATR). **A1.** light stimulation (n=199) **A2.** air puff and light stimulation (n=509) **Top**. ethogram, showing behavioral sequences of each individual (one line in the ethogram)as a function of time. **Bottom.** mean behavioral probability across the population. **B.** optogenetic activation of Basins (L72F11>CsChrimson) with R11A07 inactivated (R11A07>TNT) alone (B1, B2 or combined with air puff (B3, B4) **B1.** larvae reared without ATR, (n=45)**, B2.** with ATR (**B2** (n=48) During air-puff stimulation **B3.** without ATR (n=190), with ATR (**B4**(n=255**) C.** Optogenetic activation of Basins (L72F11>CsChrimson) alone (C1, C2 or combined with air puff (C3, C4) **C1.** Larvae reared without ATR (n=102), **C2.** with ATR (n=72). **C3. C4.** During air-puff stimulation: **C3.** larvae reared without ATR (n=147), **C4.** with ATR (n=583). Light intensity, 0.3mW/cm², average air puff intensity 4m/s.

**Supplementary figure 11.**
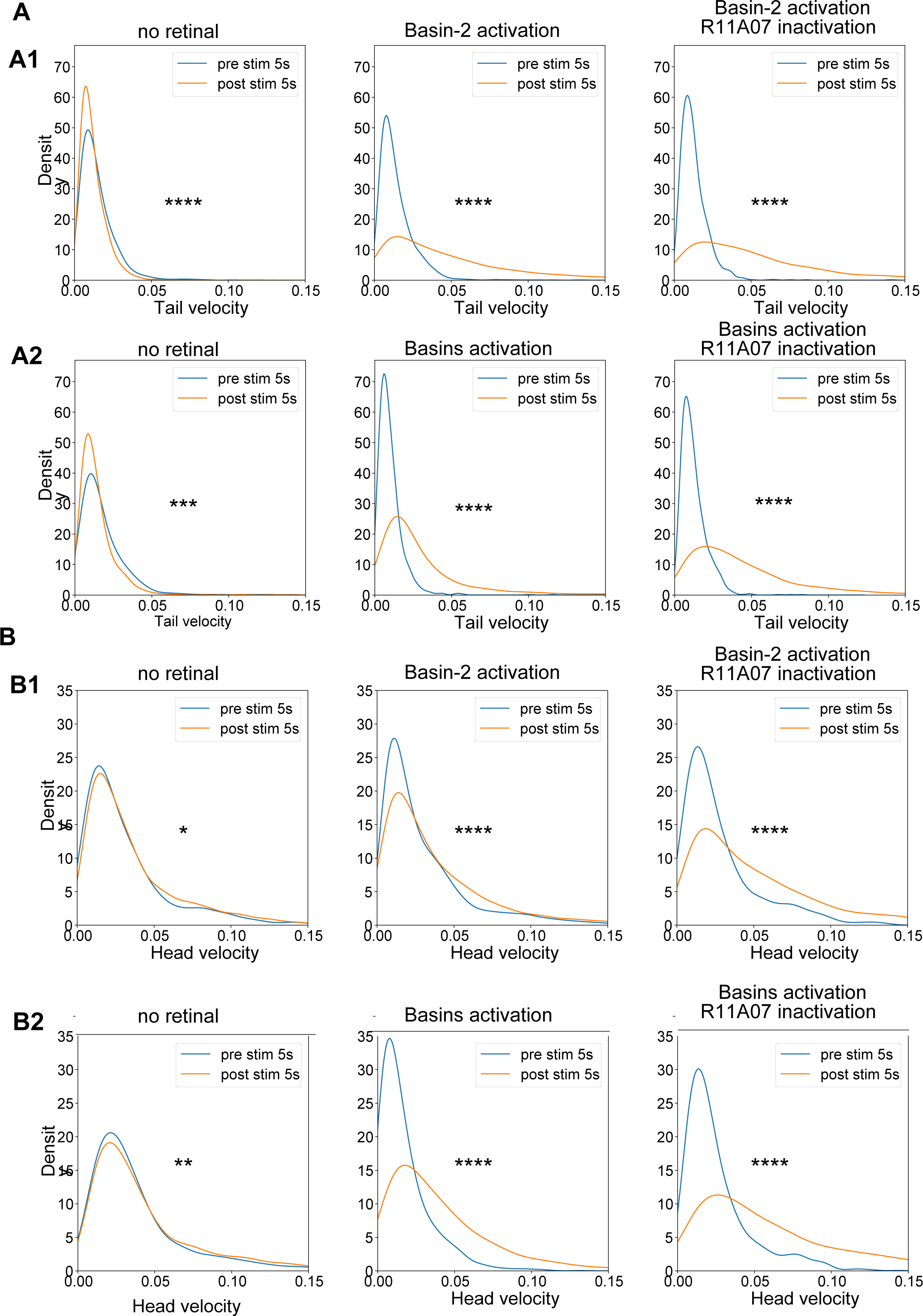
Tail and Head velocity **A.** Tail velocity (in s^−1^) (during Head cast) during the first five seconds upon stimulation compared to a same duration time window prior to stimulus onset. **A1.** Basin-2 activation: no-retinal control (left), larvae with basin-2 activated (middle) and basin-2 activated and R11A07 inactivated (right) **A2**. Basin-2 activation: no-retinal control (left), larvae with basin-2 activated (middle) and basin-2 activated and R11A07 inactivated (right. **B.** Head velocity (in s^−1^) (during Head cast) during the first five seconds upon stimulation compared to a same duration time window prior to stimulus onset.) **B1-B2.** Genotypes as in A. *:p<0.05, **:p<0.001, ***:p<0.0001, ****:p<0.00001, Kolmogorov-Smirnov test.

**Supplementary figure 12.**
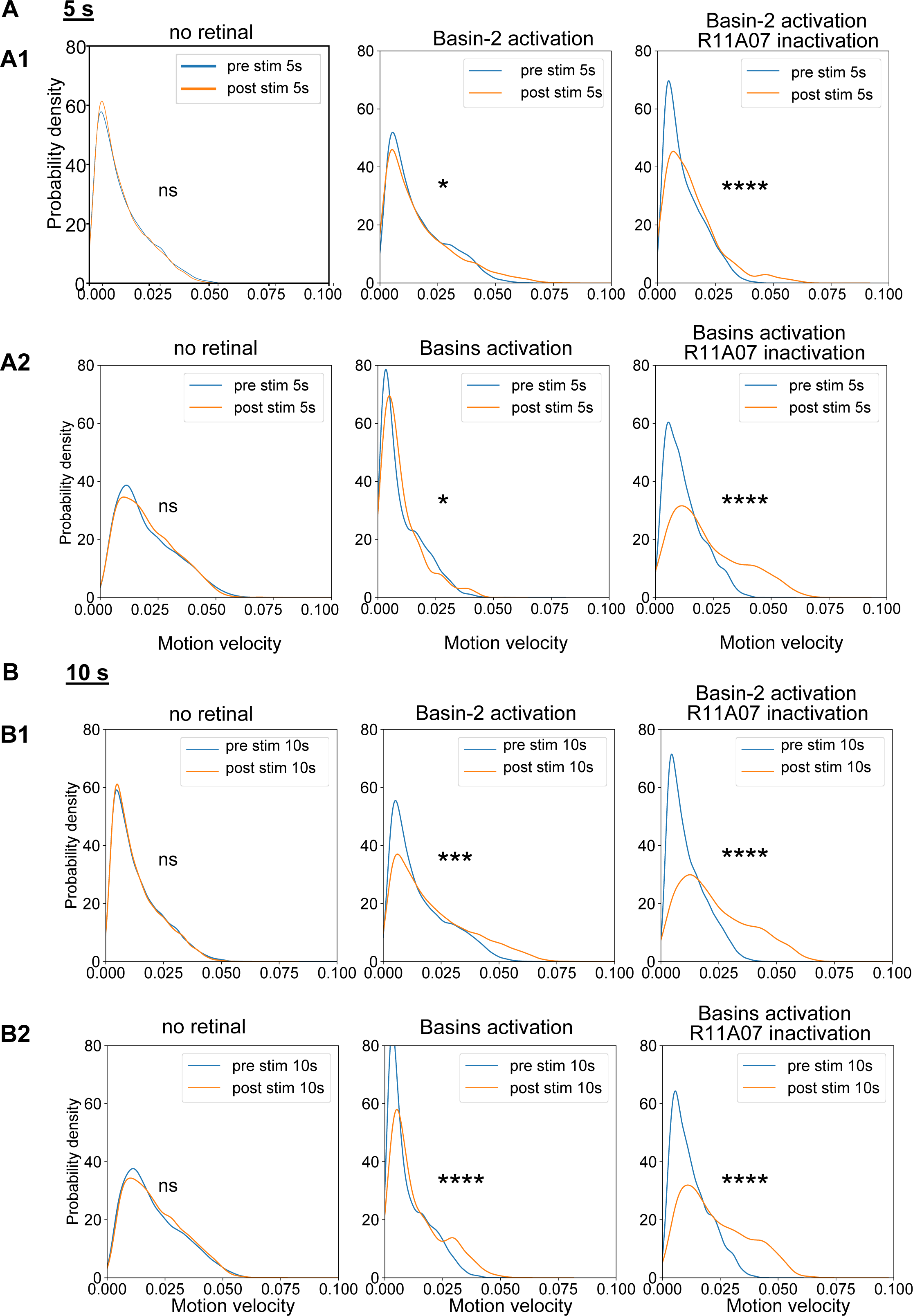
Distribution of Motion velocities **A.** Motion velocity during crawls (^-^in s^−1^) during the first five seconds upon stimulation compared to a same duration time window prior to stimulus onset. **A1.** Basin-2 activation: no-retinal control (left), larvae with basin-2 activated (middle) and basin-2 activated and R11A07 inactivated (right) **A2**. All Basins activation: no-retinal control (left), larvae with all Basins activated (middle) and all Basins activated and R11A07 inactivated (right) **B.** Motion velocity during crawls (^-^in s^−1^) during the first 10 seconds upon stimulation compared to a same duration time window prior to stimulus onset. **B1-B2.** Genotypes as in A. *:p<0.05, **:p<0.001, ***:p<0.0001, ****:p<0.00001, Kolmogorov-Smirnov test

**Supplementary figure 13.**
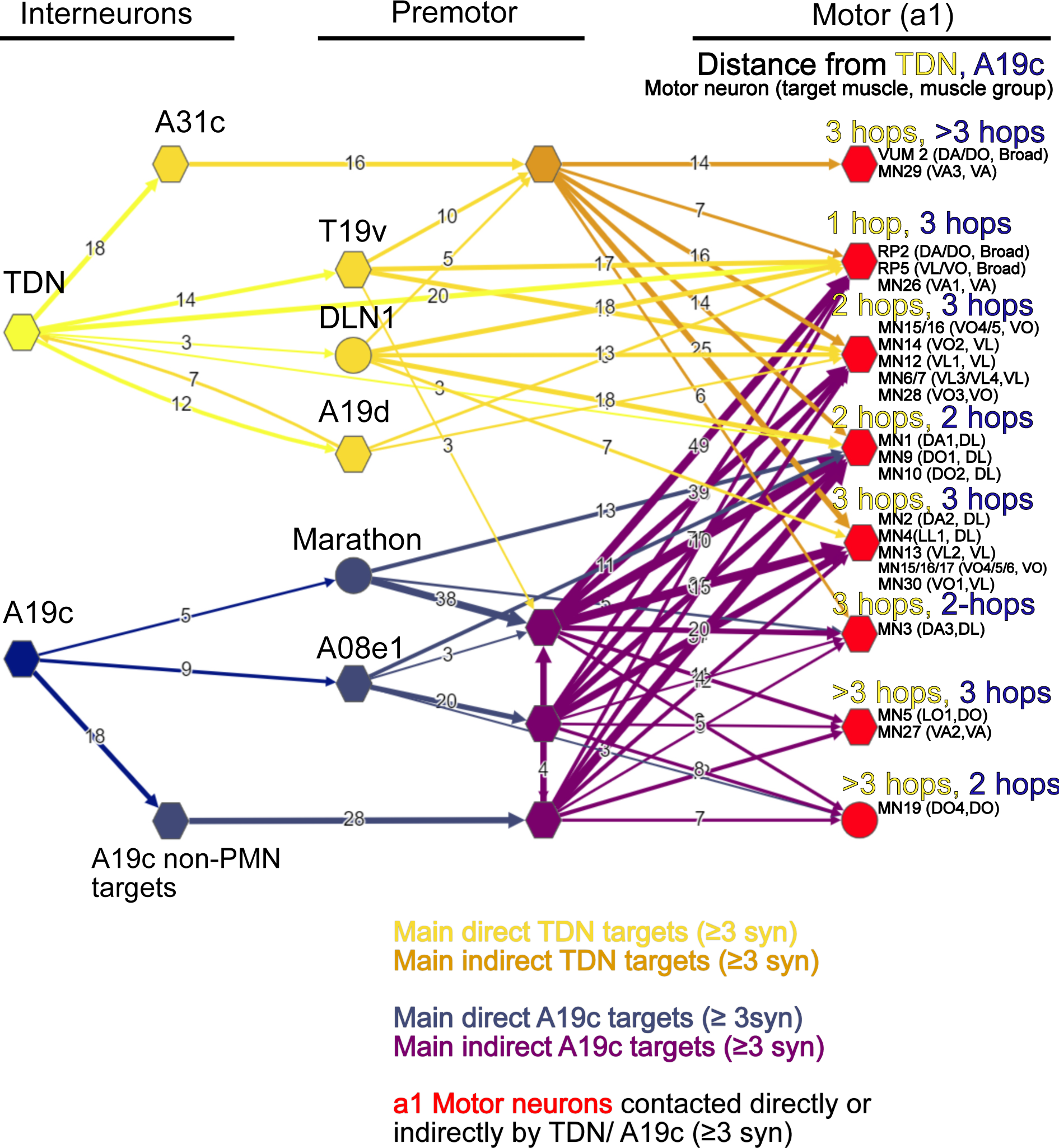
Detailed postsynaptic connectivity of TDN, A19c. Main pathways from TDN to a1 motor neurons (MNs), and from A19c to a1 MNs. Shown are all neurons receiving 3 or more synapses (≥ 3 syn) from either TDN or A19c, that in turn were significantly connected (≥ 3 syn) to either a1 MNs or to premotor neurons (indirect targets), themselves significantly connected (≥ 3 syn) to any a1 MNs. Neurons which received significant input fromTDN/A19c, but did not connect significantly to a1MNs were excluded. 22 MNs from segment a1 received significant (≥ 3 syn) direct or indirect contact from TDN and/or A19c. These 22 MNs were placed into 8 groups (red), depending on their synaptic distance to TDN and A19c. The full list of neurons shown in this graph is detailed in Supplementary table 4.

**Supplementary figure 14.**
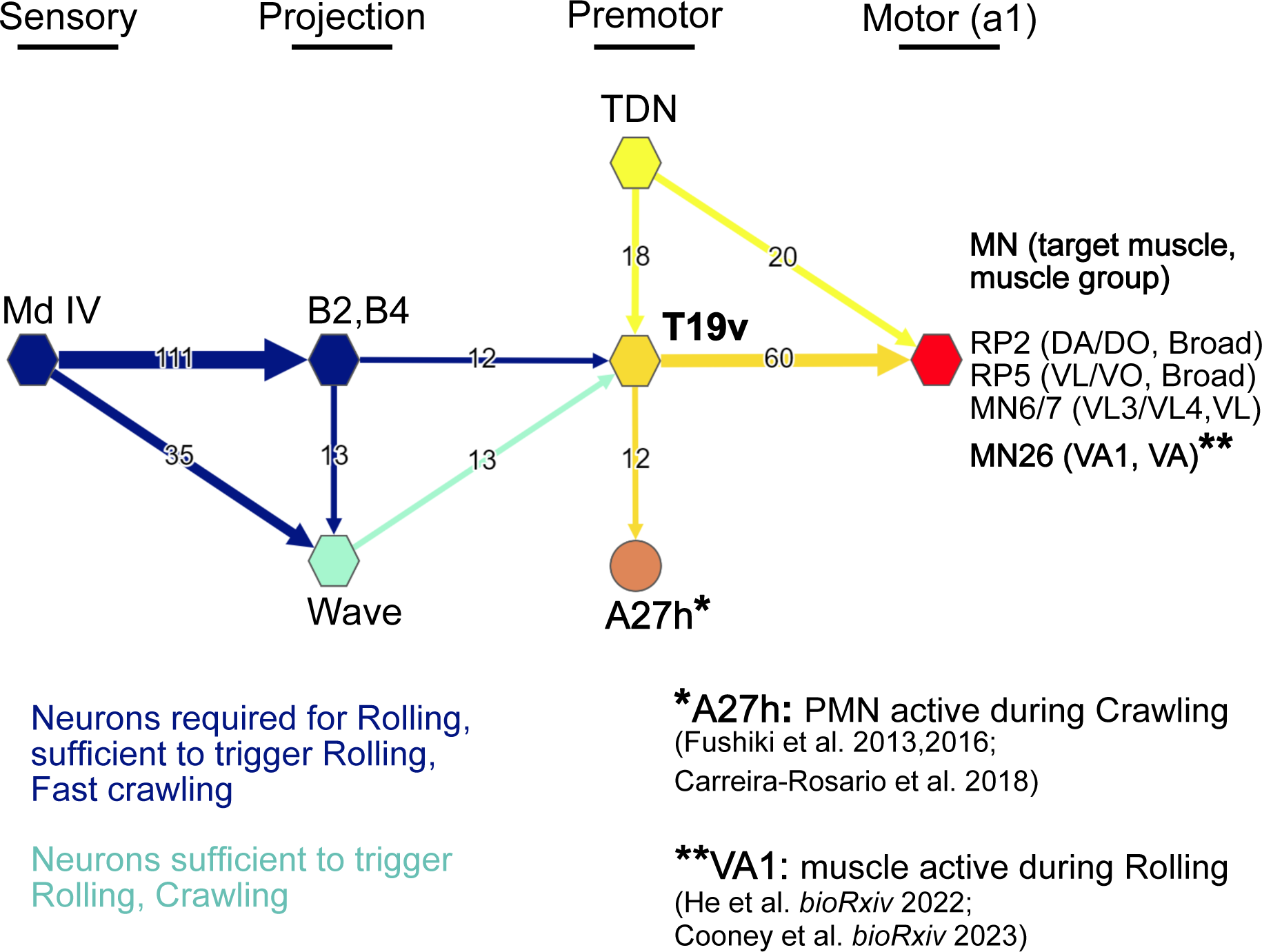
premotor neuron T19v is connected to neurons involved in Rolling and Crawling. Shown in this plot are T19v (t1,t2), its main presynaptic partners shown to be involved in Rolling: Wave (a2), Basins 2 and 4 (a1), both receiving input from Md IV nociceptive neurons (a1), TDN, which this study suggests may inhibit or promote Roll. Also shown are the main motor neurons contacted by T19v. Among them is MN26, controlling the muscle VA1, recruited during Rolling. The premotor neuron A27h (here from segment a3), shown to be significantly involved in forward Crawling, is one of the main targets of T19v. Since Rolling is often followed by Fast-Crawling, T19v position makes it an interesting candidate for the sensorimotor control of Rolling but also for the motor control of the transition from a Roll into a Fast-Crawl. It should be noted that T19v also receives significant input from Wave neurons in neuromere a1, but due to the segment-specific nature of Wave connectivity and function, only Wave from neurome a2 was included, as activation of Wave in segments a2-a6 was shown to trigger Rolling (Takagi et al. 2017). This connectivity graph was made with the web-based EM database software CATMAID.

**Supplementary atlas.**
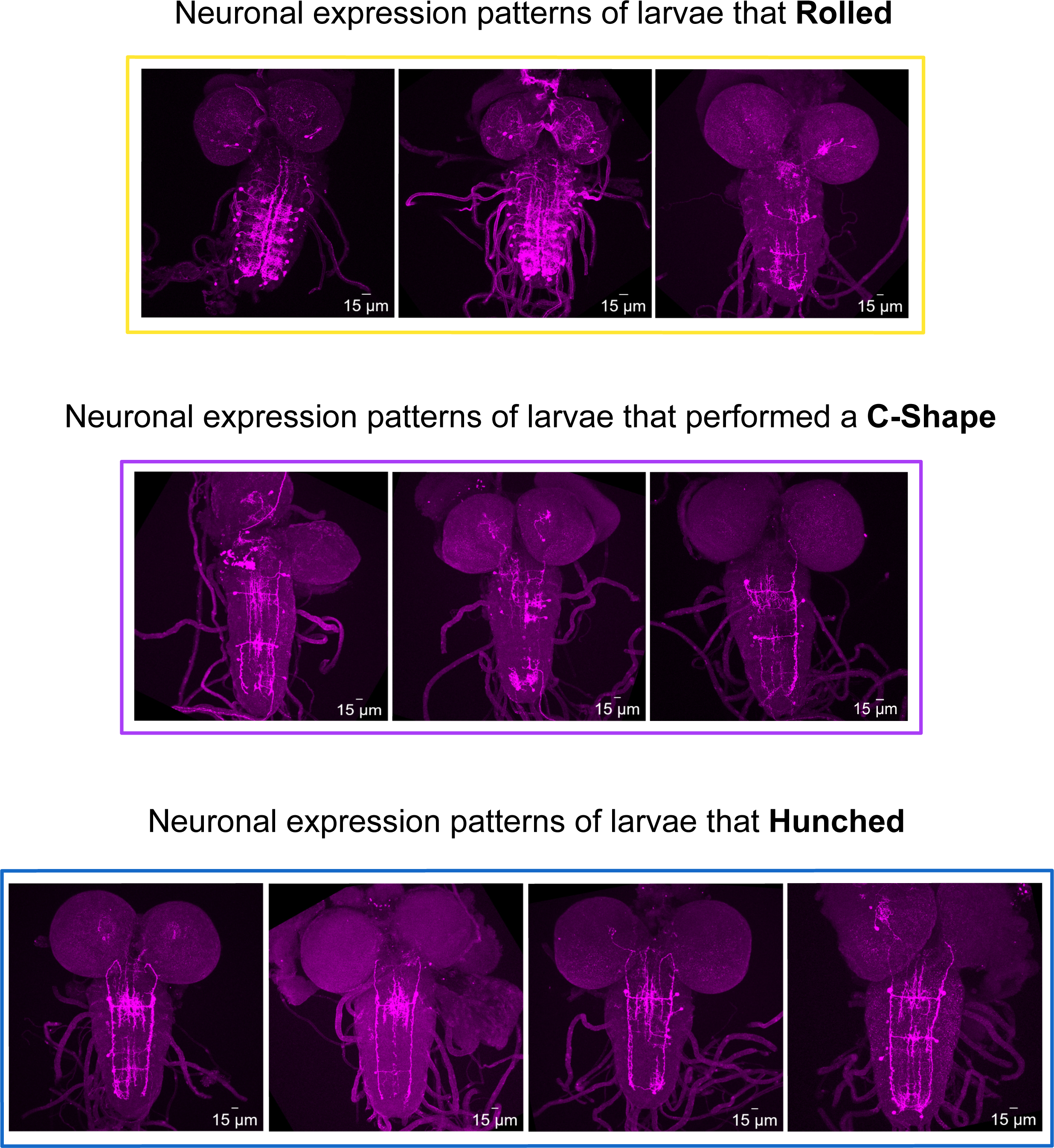
Expression profile of brains of larvae that hunched, rolled or performed a C-shape.

